# Computational Analysis of Insulin-Glucagon Signalling Network: Implications of Bistability in Metabolic Homeostasis and Disease states

**DOI:** 10.1101/509208

**Authors:** Pramod R Somvanshi, Manu Tomar, Venkatesh Kareenhalli

**Affiliations:** Department of Chemical Engineering, Indian Institute of Technology, Bombay, Powai, Mumbai, India; Bioengineering Division, John A. Paulson School of Engineering and Applied Science, Harvard University, Cambridge, USA

**Keywords:** Insulin signalling, Glucagon signalling, bistability, metabolic homeostasis

## Abstract

Insulin and glucagon control plasma macronutrient homeostasis through their signalling network composed of multiple feedback and crosstalk mechanisms. To understand how these interactions contribute to metabolic homeostasis and disease states, we analysed the steady state response of metabolic regulation (catabolic or anabolic) with respect to structural and input perturbations in the integrated signalling network, for varying levels of plasma glucose. Structural perturbations revealed: the positive feedback of AKT on IRS is responsible for the bistability in anabolic zone (glucose >5.5 mmol); the positive feedback of calcium on cAMP is responsible for ensuring ultrasensitive response in catabolic zone (glucose <4.5 mmol); the crosstalk between AKT and PDE3 is responsible for efficient catabolic response under low glucose condition; the crosstalk between DAG and PKC regulates the span of anabolic bistable region with respect to plasma glucose levels. The macronutrient perturbations revealed: varying plasma amino acids and fatty acids from normal to high levels gradually shifted the bistable response towards higher glucose range eventually making the response catabolic or unresponsive to increasing glucose levels. The analysis reveals that certain macronutrient composition may be more conducive to homeostasis than others. The network perturbations that may contribute to disease states such as diabetes, obesity and cancer are discussed.

## Introduction

Living systems deploy bio-molecular networks comprising of multiple feedback loops to facilitate an optimal response to an environmental stimulus. The network structure and its kinetics define steady state and dynamical properties of the output response^1^. One such property is homeostasis, wherein the levels of physiological variables are held in a narrow range despite any external perturbation to the system^2^. One of the optimal strategy to obtain homeostasis is to have bistable control in the regulatory circuit ^3^. Bistability is a property in which the threshold for activation and deactivation of the response differs (hysteresis) leading to two stable states for a given stimulus, depending upon the history of the stimulus ^4^. Several biological systems exhibit bistability such as cell cycle ^5^, MAPK cascade and JNK signalling ^6,7^, immune response ^8^, insulin signalling pathway ^9^ and neurological states^10^. Bistable circuits are known to impart switch like response, robustness to noise, memory of the stimulus, and irreversibility in response^11-13^. Disturbances in the operation of bistable response is implicated in dysregulation of homeostasis and subsequent disease states such as diabetes, obesity and cancer ^14,15^. Bistability with respect to PKC response in insulin signalling network could explain selective hepatic insulin resistance ^16^. Bistability in AKT response with respect to insulin levels is also reported through simulations of insulin/AKT and MAPK/ERK signalling pathways^14,17,18^. Furthermore, for the physiological range of plasma glucose levels, the flux through glycolysis exhibits multiple steady state responses in HeLa cells ^19^ indicating the interplay between the regulatory feedback loops and their effector hormonal signals. It is known that these pathways are regulated by insulin and glucagon, motivating further analysis on underlying complexity of hormonal regulation of metabolism ^20^. Therefore, in this study we focus on analysing the effects of perturbations in the network structure and multiple-input stimulus on the response of insulin-glucagon signalling network and its relation to metabolic homeostasis and disease states.

While insulin is an anabolic hormone, glucagon is a catabolic hormone. Both modulate each other to maintain the level of key metabolites like glucose, fatty acids and amino acids in plasma ^21,22^ under different physiological conditions (resting, postprandial and exercise). These hormones act antagonistically towards each other at the stages of their secretion and signalling (See supplementary file Appendix I). Insulin not only stimulates glucose uptake and lipid synthesis but also inhibits lipolysis, proteolysis, glycogenolysis, gluconeogenesis and ketogenesis in tissues like liver, muscle & adipose^23,24^. Glucagon, on the other hand, mediates catabolic pathways and renders elevation in levels of plasma metabolites like fatty acid, glucose and amino acids in order to supply body’s physiological needs ^25^ under relevant conditions. Further, plasma macronutrient are known to regulate the secretion and signalling of insulin and glucagon. Glucose is known to induce insulin secretion and inhibit glucagon secretion ^26,27^. Amino acids induce both insulin and glucagon secretion in a threshold dependent manner^28,29^. While amino acids activate insulin signalling through AKTp ^30^, it inhibits IRS through S6kp activation ^31^. Fatty acids can induce insulin secretion and inhibit insulin signalling at higher plasma levels^32,33^. These varied interaction of the macronutrients with the hormonal regulatory mechanisms results in a highly nonlinear regulatory response for different combinations of these macronutrients in the plasma. Therefore, it is interesting to study how the bistability in the insulin-glucagon network and resultant metabolic state varies with different macronutrient compositions.

Several mathematical model of insulin signalling pathways have already been documented^34-36^. Moreover, subsystem models of insulin receptor binding, receptor recycling & GLUT4 translocation leading to glucose uptake is also included in the overall insulin signalling model ^37^. Mathematical models explaining signal transduction by G protein and downstream calcium signalling have been proposed in literature^38-40^. These models explain the dynamics of G-protein activation and receptor desensitisation based on ligand binding and subsequent events involving calcium and PLC^41-43^. Although several models differently look at insulin and glucagon signalling and its regulation of metabolism, there is scarcity of models that consider the mutually antagonistic effect of insulin and glucagon signalling and the effect of macronutrients on the interplay of these pathways. Therefore, we have developed and analyzed an integrated model of insulin-glucagon signalling to obtain insights on the metabolic response with respect to different levels of glucose, amino-acid and fatty acid in the plasma (see Figure 1). In the current study, we analyse the effect of varying levels of plasma macronutrients and knock-out of feedback loops and crosstalk mechanisms on phosphorylation state elicited by the integrated network. We also report the bistable response in the network over certain range of physiological conditions and its disruption indicating disease state.

**Figure 1.**
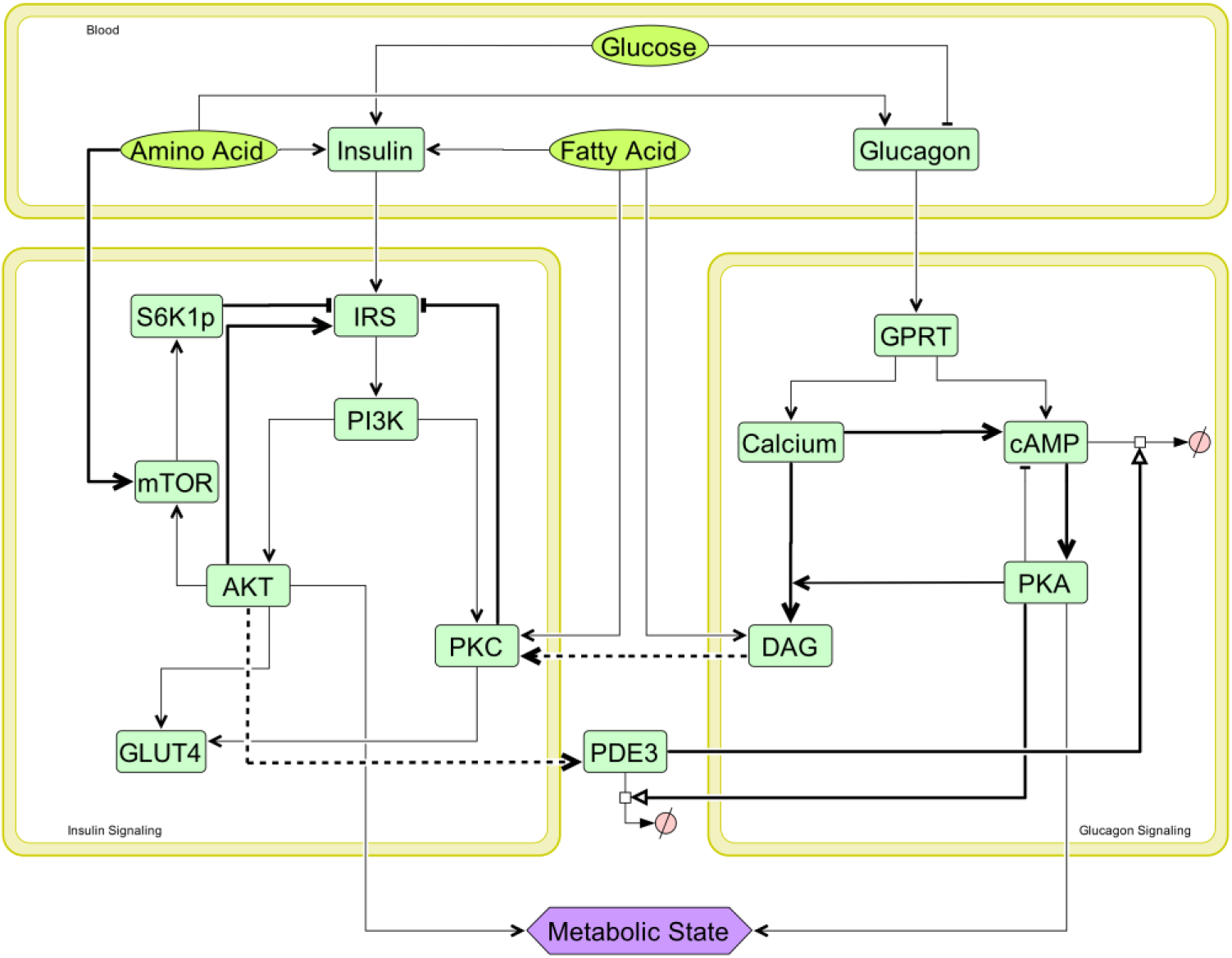
Integrated insulin-glucagon signalling network. Bold lines represent positive feedbacks (→), negative feedbacks (—|) and crosstalks (→) in the network. There are three modules in the network, namely - insulin signalling, glucagon signalling and blood. Glucose, amino acids & fatty acids are the input macronutrients present in plasma. Based on the amount of these macronutrients in different physiological situations, pancreas secrete different amounts insulin and glucagon in plasma. These hormones then trigger corresponding signalling pathways in tissues like liver, fat and muscle. Insulin and glucagon signalling modules act antagonistically to each other with the help of crosstalk (AKTp activating PDE3 which promotes cAMP degradation in glucagon signalling; and DAG activates PKC which inhibits IRS in insulin signalling) and feedbacks. When insulin signalling fluxes are greater than glucagon signalling fluxes, net metabolic state is anabolic; and when glucagon signalling fluxes are greater than insulin signalling fluxes, net metabolic state is catabolic.

## Results

The mathematical model for insulin-glucagon integrated network was simulated to obtain the steady state profiles of activated AKT, PKA and the phosphorylation state (*P*_*s*_) for various levels of plasma glucose concentration. The steady state profiles were obtained for both switching ON (i.e., increasing glucose levels) and switching OFF (i.e., decreasing glucose levels). The phosphorylated AKT (AKTp) profile show a typical bi-stable response, with the activation of AKTp occurring at 1.6 fold of physiological glucose levels (5mM) and deactivation at about 1.2 fold (Figure 2 (i)). In contrast, the activation of PKA is monostable, with a highly sensitive response (Hill coefficient = 5.8, Figure 2(ii)).

**Figure 2.**
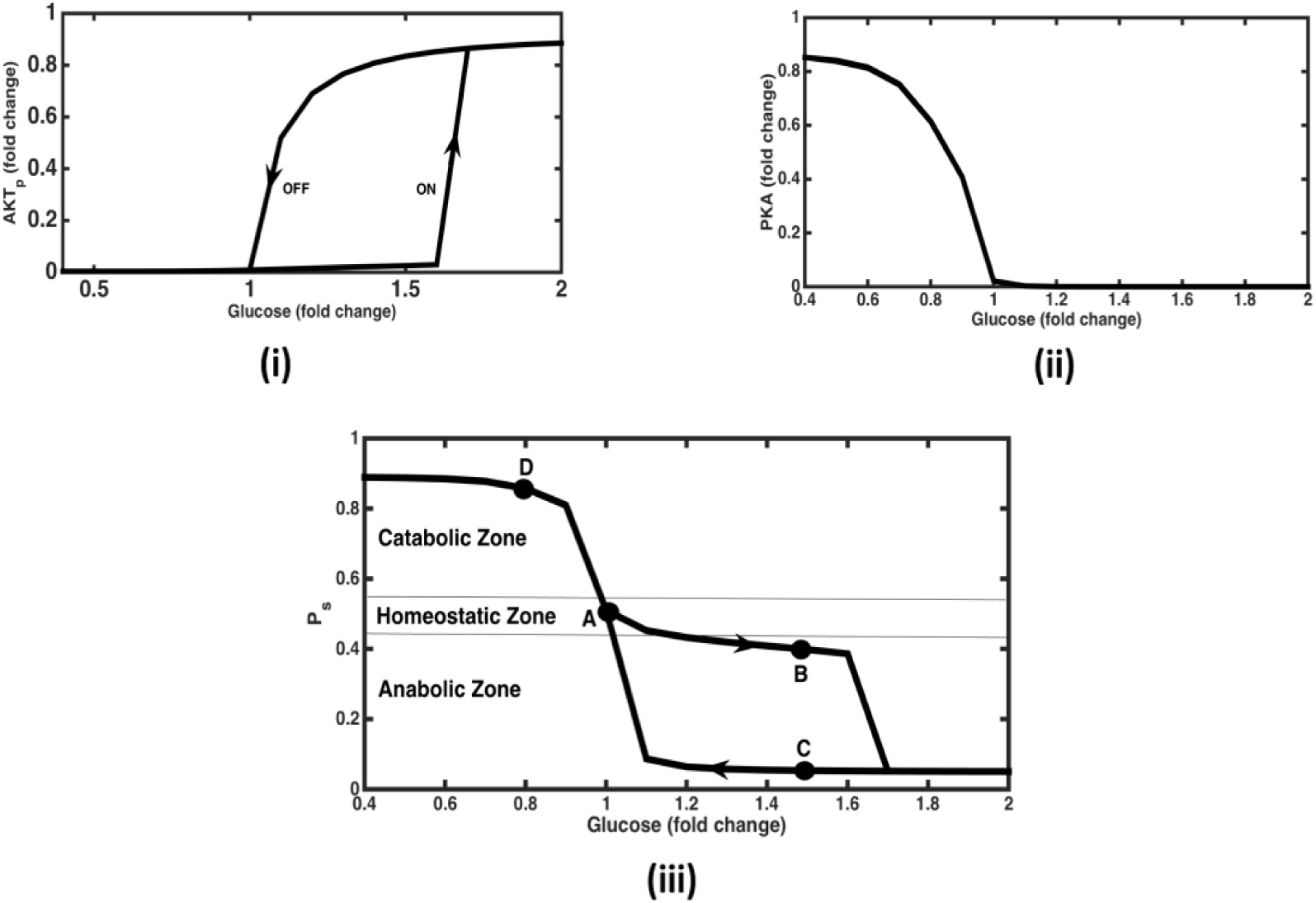
The steady state response for AKTp, PKA and *P*_*s*_ (Phosphorylation state) with varying levels of glucose in plasma. (i) AKTp vs Glucose response is bistable and ultrasensitive. Response increases suddenly at glucose = 1.6 folds while switching ON (increasing levels of glucose, denoted by the arrow pointed upwards) and decreases to almost zero level at glucose = 1fold while switching OFF (decreasing levels of glucose, denoted by the arrow pointing downwards). (ii) PKA vs Glucose response is monostable and ultrasensitive. PKA level drops sharply to almost zero for glucose levels beyond 1fold. There no difference in switching ON and switching OFF paths. (iii) *P*_*s*_ vs Glucose response. Here bistability in AKTp response is getting translated in *P*_*s*_ (a combination of AKTp and PKA response, see equation 1) response. *P*_*s*_ levels are greater than 0.51 for glucose levels less than 1fold, tending towards catabolic response. *P*_*s*_ levels remain in the homeostatic range for glucose ranging between 1 to 1.6 folds and then turn anabolic beyond glucose levels of 1.7 folds (while switching ON). While switching OFF, *P*_*s*_ levels remain anabolic up to 1.1 folds and then turn homeostatic and catabolic at lesser levels of glucose. Note that *P*_*s*_ levels greater than 0.55 indicates catabolic response, between 0.44 and 0.55 indicates homeostatic response and less than 0.45 indicates anabolic response. Points A, B, C & D denote different steady state conditions. A – Glucose, *P*_*s*_ = 0.51 (homeostasis). B - Glucose = 1.5 folds (switching ON), *P*_*s*_= 0.4 (mildly anabolic). C – Glucose = 1.5 folds (switching OFF), *P*_*s*_ = 0.05 (highly anabolic). D – Glucose = 0.8 fold, *P*_*s*_ = 0.86 (highly catabolic).

To characterise the metabolic state under different perturbations for varying glucose levels, *P*_*s*_ was determined and is shown in Figure 2 (iii). It demonstrates a bistable response, however only in the lower half of the phosphorylation state representing the anabolic zone due to AKT phosphorylation. Under resting state, at a glucose concentration of 5 mM, phosphorylation state value is equal to 0.51 showing slightly catabolic conditions under fasting condition. Under conditions of increased glucose consumption (such as during physical activity, stress etc.) with decrease in levels of plasma glucose, the *P*_*s*_ value attains greater than 0.5 indicating a catabolic state without any bistability (also reflected by the absence of bistability in the PKA vs glucose response (Figure 2 (ii)).

On increasing glucose concentration from the fasting state, the *P*_*s*_ transits through a mildly anabolic phase, with a dedicated fully operational anabolism occurring at higher glucose concentration (1.7 fold change). This implies that for glucose perturbation of less than 1.7 fold, *P*_*s*_ remains only mildly anabolic. On perturbing glucose levels greater than 1.7 fold, *P*_*s*_ is highly anabolic and remains in this state even after lowering of glucose up to 1.1 fold of the resting state. This indicates that the anabolic fluxes such as glycogenesis and lipogenesis, if increased beyond certain threshold, might get locked even after lowering the glucose concentration, due to the bistability in *P*_*s*_ with respect to glucose.

It is interesting to note that a buffering zone exists for the anabolic response, while such a bistable response is absent in the catabolic zone. Glucose concentration lower than the physiological levels switches *P*_*s*_ value closer to one indicating mainly catabolism. Thus, the bistability offers a buffering zone between 1 to 1.7 fold change of glucose, in which both mainly anabolic and a balance between anabolic and catabolic process can be observed for a given glucose concentration, depending on the ON and OFF path. This indicates that for a given glucose concentration two distinct metabolic states can be achieved depending on the steady state points on the two different paths.

In order to elaborate on the operating strengths of feedbacks and crosstalk conditions in different steady states (shown by alphabets A-D in Figure 2(iii), we have shown feedback strengths operating in the network (see Figures S2 – S5). Under fasting conditions (represented by point A in Figure 2 (iii)), Figure S2 shows that the dominant feedback is PDE3 degradation of cAMP thereby inhibiting the catabolic module. Hence the positive feedbacks of Ca on DAG & cAMP along with cAMP on PKA are almost inactive. Further, anabolic module is also minimally active due to low levels of insulin secretion under these conditions leading to minimal activation of the positive feedback loop involving IRS-PI3K-AKT. When glucose levels increase to 1.5 folds (shown as point B in Figure 2 (iii)), insulin secretion goes up thereby activating the anabolic module (Figure S3). In this case, increased degradation of cAMP by AKT (via PDE3 crosstalk) leads to complete shut off of the catabolic regime. In insulin signalling module, AKT positive feedback on IRS increases slightly due to increased insulin secretion, but is not enough to make *P*_*s*_ heavily anabolic. Note that the feedbacks and crosstalk like DAG inhibiting IRS via PKC and amino acids inhibiting IRS via mTOR-S6K signalling are almost inactive under these conditions.

On the other hand, under switching OFF conditions (shown as point C in Figure 2(iii)), the positive feedback of AKT on IRS increases along with the PKC and mTOR inhibitions of IRS (Figure S4). But the overall effect is dominated by the positive feedback loop between IRS-PI3K-AKTp, thereby locking the state in highly anabolic region. Further, PDE3 degradation of cAMP is highly active causing the complete inactivation of glucagon signalling (just like the state B). Whereas under low glucose conditions (shown as point D in Figure 2(iii)), anabolic module becomes inactive due to low levels of insulin secretion from pancreas leading to reduced degradation of cAMP due to AKT via PDE3 creating a highly catabolic state (Figure S5). Further, increased PKA activates DAG which inhibits IRS through crosstalk with PKC leading to inactivation of anabolic signalling pathways.

### Effect of varying plasma amino acid levels

In order to assess the effect of macronutrients on *P*_*s*_ with respect to glucose, the level of fatty acids (FA) and amino acids (AA) in plasma were varied. When AA level was increased up to 3 folds (while keeping FA constant at 1 fold), the span of bistable response increased in the anabolic region (Figure 3(i)). The *P*_*s*_ response now turns anabolic at glucose = 1.8 folds, (vis-a-vis at glucose = 1.7 folds when AA = 1 fold) while switching ON and turned homeostatic (from anabolic) at glucose = 0.9 fold (vis-a-vis glucose = 1 fold when amino acid = 1 fold) while switching OFF. Corresponding network map (Figure S6, representing point E in Figure 3(i)) shows high activation of insulin signalling module due to dominance of AKTp positive feedback on IRS under such conditions. It also leads to increased cAMP degradation by PDE3 and thus supresses the effect of glucagon signalling module. This is why overall metabolic state is heavily anabolic in spite of glucagon signalling module also being activated due to increased glucagon secretion under high levels of AA.

**Figure 3.**
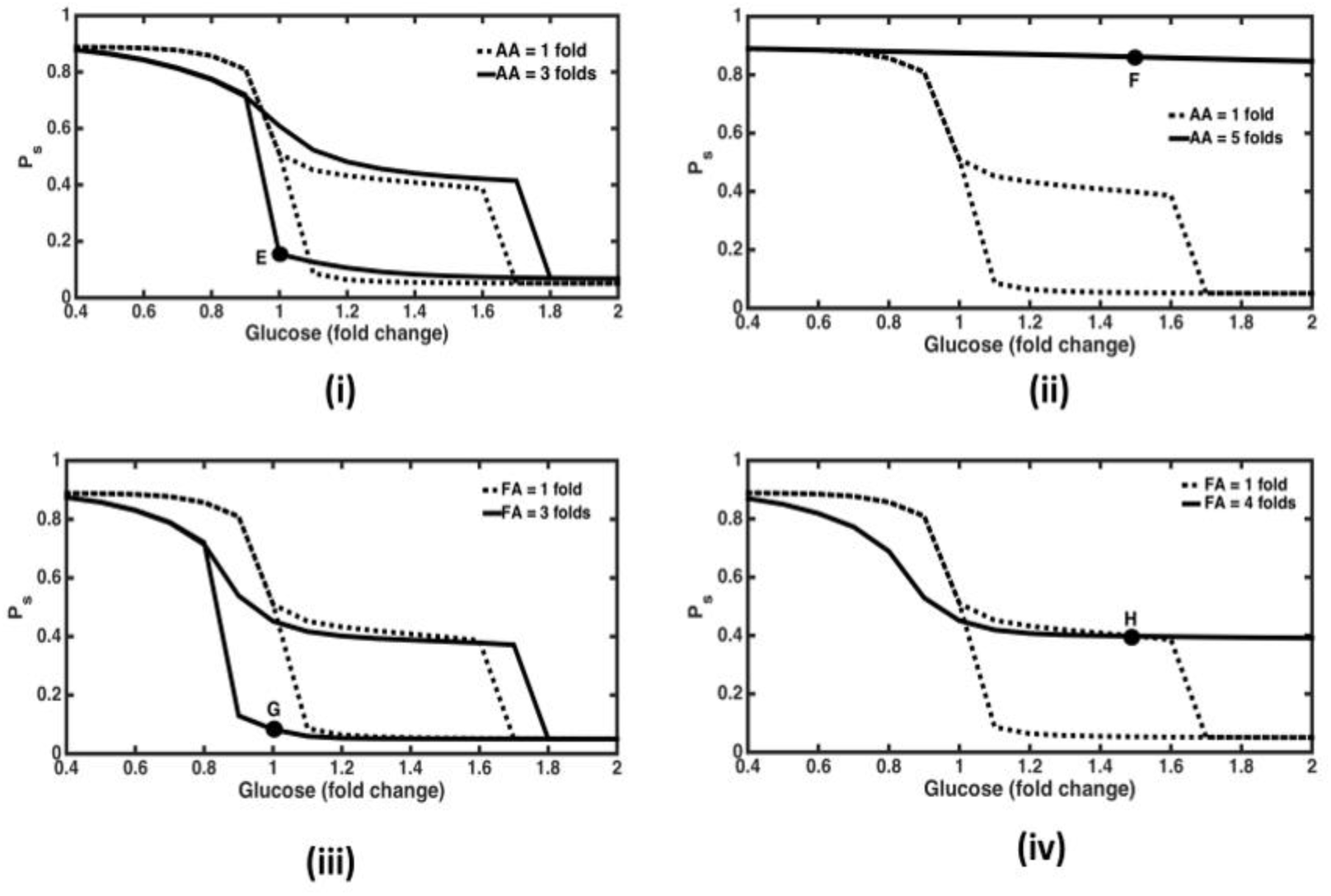
Effect of changing macronutrients on bistability. Dashed graphs show normal responses whereas solid graphs show responses at higher levels of macronutrients (amino acid and fatty acid) levels in plasma. (i) AA = 3 folds. Here bistability span is increasing in the anabolic zone with response turning anabolic at 1.8 folds (compared to 1.7 folds for AA = 1 fold) while switching ON, and response turning catabolic from anabolic at 0.9 fold (compared to 1 fold for AA = 1 fold, where *P*_*s*_ turns homeostatic from anabolic) while switching OFF. At point ‘E’ glucose = 1 fold, *P*_*s*_ = 0.15 and corresponding anabolic conditions are depicted in Figure S5. (ii) AA = 5 folds. Here response remains catabolic even at glucose levels up to 2 folds. At point ‘F’ glucose = 1.5 folds with *P*_*s*_ = 0.86, corresponding network diagram is shown in Figure S6. (iii) FA = 3 folds. Here again the span of bistable response is increasing in the anabolic space with *P*_*s*_ value turning anabolic at 1.8 folds (as compared to 1.7 folds for FA = 1 fold) while switching ON, and response turning catabolic from anabolic at glucose = 0.8 fold (as compared to glucose = 1 fold for FA = 1 fold, where *P*_*s*_ turns homeostatic from anabolic) while switching OFF. At point ‘G’, glucose = 1 fold and *P*_*s*_ = 0.08; corresponding network diagram is shown in Figure S7. (iv) FA = 5 folds. Here response remains monostable and non-anabolic even at high glucose levels up to 2 folds. *P*_*s*_ turns homeostatic (from catabolic) at subnormal glucose levels of 0.9 fold (as compared with normal glucose levels = 1 fold for FA = 1fold). At point ‘H’, glucose = 1.5 folds, *P*_*s*_ = 0.39; corresponding network representing mildly anabolic conditions are shown in Figure S8.

On further increasing AA to 5 folds, *P*_*s*_ response turns catabolic and monostable even at high glucose levels up to 2 folds. *P*_*s*_ remains above 0.9 in this case (Figure 3(ii)). Here active AKT levels are always low with sub-sensitive and higher levels of active PKA response with respect to glucose, resulting in a broad catabolic state. This indicates an abnormal physiological state wherein the anabolic pathways are not activated even at high levels of plasma glucose. Corresponding network map (Figure S7 representing point F in Figure 3(ii)) shows high levels of mTOR activation by AA ultimately leading to increased inhibition of IRS by S6K1p ^44^. This inactivation of insulin signalling leads to reduced inhibition of PDE3 by cAMP. Further, high AA levels also increase glucagon secretion from pancreas ^45^ creating a highly catabolic state in the network.

### Effect of varying plasma fatty acid levels

Similarly, on increasing FA levels up to 3 folds, while keeping AA constant at 1fold, we again observe an increase in the span of bistable response (Figure 3(iii)). The response now turns anabolic at glucose = 1.8 folds (vis-a-vis at glucose = 1.7 folds when FA = 1 fold) while switching ON, and the response turns homeostatic (from anabolic) at glucose = 0.8 folds (vis-a-vis at glucose = 1 fold when FA = 1 fold) while switching OFF. Corresponding flux map (Figure S8, representing point G in Figure 3(iii)) depicts high levels of IRS activation mainly due to the positive feedback of AKT which is keeping the overall metabolic state as anabolic in spite of inhibition by increased FA levels (via PKC). Moreover high levels of activation of PDE3 degradation of cAMP is also shutting off the catabolic signalling module under these conditions.

On further increasing FA levels to 5 folds, *P*_*s*_ response becomes monostable and does not turn anabolic even at glucose levels up to 2 folds (Figure 3(iv)). Here AKTp levels are very low and PKA response is hyper sensitive and becomes close to zero as glucose levels go above 1 fold leading to overall homeostatic response at higher glucose levels in this case. Corresponding flux map (Figure S9, representing point H in Figure 3(iv)) shows that insulin signalling is inhibited by higher FA levels (via PKC activation) ^46^ even at high glucose levels. Moreover, glucagon signalling is inactivated due to inhibition of glucagon secretion by glucose and significant level of cAMP degradation by PDE3.

### Knockout of positive feedback of AKT on IRS

In order to evaluate the effect of feedbacks on the *P*_*s*_ response, the positive feedback from AKTp on IRS was negated. This resulted in a monostable response without the activation of anabolic response i.e., AKT was not activated (Figure 4(i)). Thus, the model suggests that this positive feedback is a dominant mechanism responsible for the anabolic bistable response. The network diagram in the absence of this feedback with glucose levels = 1.5 folds (Figure S10, representing point I in Figure 4(i)), shows that insulin signalling module is less activated (AKT contribution to *P*_*s*_ = 0.12) as compared to the corresponding conditions in the presence of this feedback during switching OFF conditions (Figure S3, where AKT contribution to *P*_*s*_ is 0.89), when the metabolic state is heavily anabolic. Here glucagon signalling module is almost shut off mainly due to highly active PDE3 degradation of cAMP and glucose inhibition on glucagon signalling. Hence due to the absence of this feedback overall metabolic response remains only slightly anabolic even at high glucose levels.

**Figure 4.**
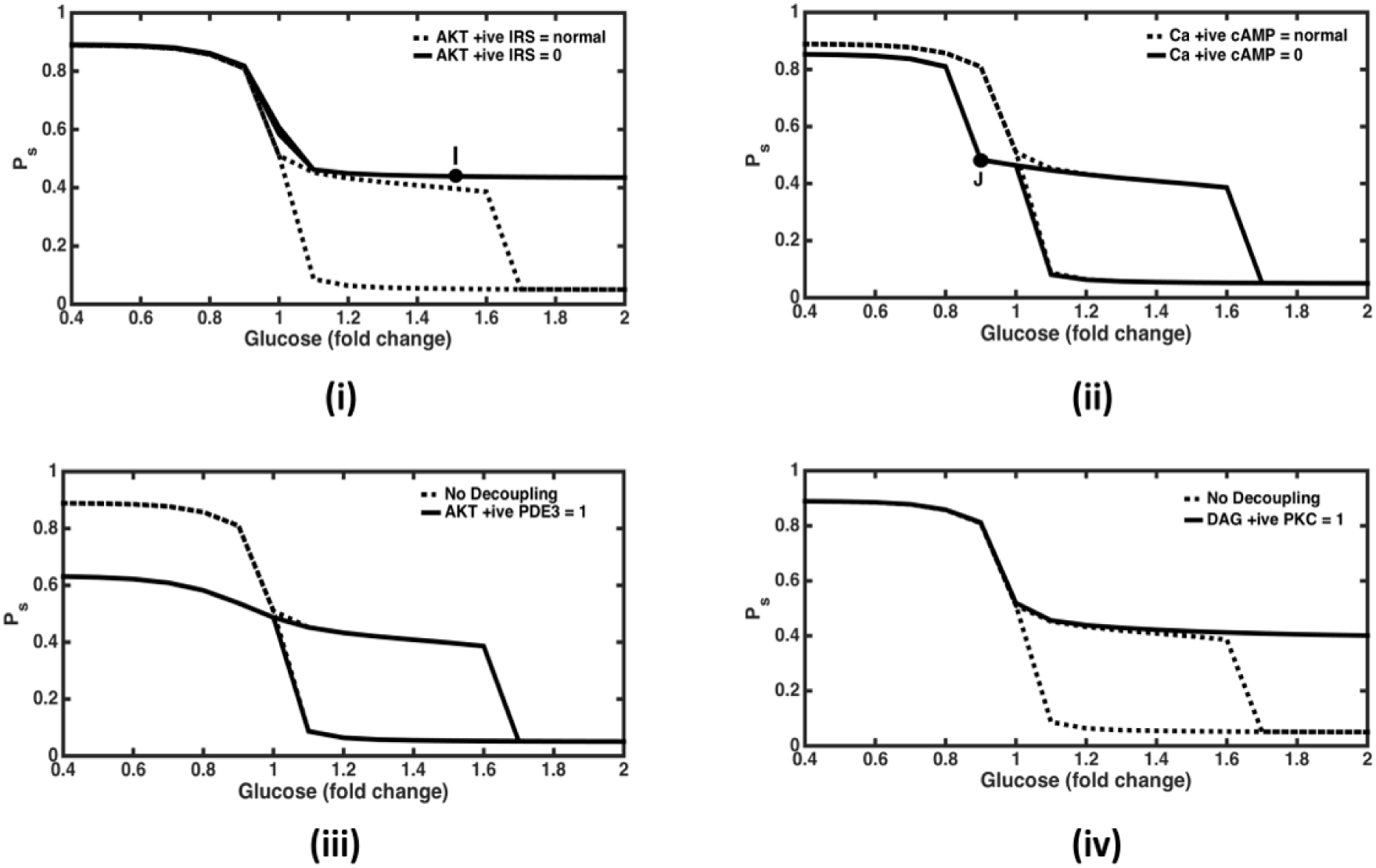
Effect of feedback and crosstalk perturbation in the network. (i) Knock out AKTp positive feedback on IRS. The response remains non-anabolic and monostable for glucose levels up to 2 folds. At point ‘I’ glucose = 1.5 folds and *P*_*s*_ = 0.44; corresponding network diagram is shown in Figure S9. (ii) Knock out of Calcium positive feedback on cAMP. Catabolic response is weakening slightly at subnormal glucose levels such that it becomes homeostatic at glucose = 0.9 fold (at ‘J’ where *P*_*s*_ = 0.48; corresponding network diagram is shown in Figure S10). (C) Decoupling of AKTp crosstalk PDE3. This leads to inactivation of glucagon signalling module at subnormal glucose levels. (D) Decoupling of DAG crosstalk with PKC. This leads to inhibition of insulin signalling module at high glucose levels up to 2 folds. This shows that the effects of both crosstalk are mutually independent, with AKTp crosstalk PDE3 playing role at subnormal glucose levels and DAG crosstalk PKC playing role at high glucose levels.

### Knockout of calcium positive feedback on cAMP

The positive feedback of calcium on cAMP was also negated to see the response in the catabolic module. It resulted in leftwards shift in the deactivation threshold of catabolic response with respect to glucose (Figure 4(ii)). This implies that it activates cAMP, PKA and DAG at subnormal glucose levels. Corresponding network diagram under these conditions (Figure S11, representing point J in Figure 4(ii)) shows how the cAMP activation of PKA and PKA contribution to the overall metabolic state is reduced as compared to the case when this positive feedback is present (Figure S5). Thus keeping the overall response as homeostatic even at subnormal glucose levels.

When both the above mentioned positive feedbacks are simultaneously knocked out, monostable and catabolic response prevails at subnormal glucose levels and monostable and homeostatic response is observed at higher glucose levels (Figure not shown). This shows that the effect of both these feedbacks is mutually independent and can be observed in different range of glucose levels.

### Knockout of crosstalk between AKT and PDE3

Next, we knockout crosstalk between insulin and glucagon signalling pathways. Firstly, we delink the activation of PDE3 by AKT. This was done by setting the Hill function representing activation of PDE3 by AKTp equal to its maximum value of 1. This resulted in a suppressed levels of *P*_*s*_ state indicating reduced activity in the catabolic zone at glucose levels less than 1 fold (Figure 4(iii)). This is mainly due to enhanced degradation of cAMP by PDE3 which results in lowered PKA response. Thus, the crosstalk helps in efficient catabolic response under low glucose conditions.

### Knockout of crosstalk between DAG and PKC

Likewise, on eliminating the crosstalk from DAG to PKC by setting the corresponding Hill function at its maximum value of 1, response turns monostable and homeostatic even at higher glucose levels up to 3 folds (Figure 4(iv)). This implies that the activation of AKTp is reduced by inhibition of IRS by this crosstalk. At higher glucose concentration, when the glucagon signalling pathway is suppressed, the DAG mediated insulin signalling suppression essentially happens due to fatty acids^47^. Therefore, the span of bistability under normal macronutrient conditions (FA = AA = 1fold) is set by the FA activation of DAG.

Overall, this shows that both the crosstalk from DAG to PKC and AKT to PDE3 are operational and relevant for normal catabolic to anabolic transitions. Further, the effect of simultaneous decoupling of both the crosstalk is algebraic summation of removal of crosstalk one at a time. This demonstrates that the effect of both the crosstalk on the integrated network response is independent of each other.

## Discussion

### Structure and function underlying bistability in IG network

The analysis of the network shows that mutually antagonistic modules of insulin & glucagon signalling increase the operational efficiencies of respective pathways for glucose as input stimulus ^3^. The analysis reveals that the robustness of a homeostatic regulatory circuit is obtained by a combination of positive and negative feedback in the system ^48,49^. While increasing the negative feedback reduces the effect of positive feedback thereby increasing the demand for a higher input stimulus; decreasing the negative feedback enhances the effect of positive feedback thereby inducing a bistable response in such a circuit ^50^. Therefore, the span of the bistable response is determined by the relative strengths of the positive and negative feedbacks in the network.

The positive feedbacks of AKT on IRS and Ca on cAMP in the pathway ensure a proportionately higher output response even for a smaller value of the input stimulus from insulin and glucagon secreted from the pancreas. On the other hand, negative feedbacks like PDE3 induced cAMP degradation, PKC & S6K inhibition on IRS are essential to reduce the effect of the input stimulus on the output response and serves as the crosstalk point for antagonistic pathways. Further, the DAG & AKT crosstalk on PKC & PDE3, respectively ensure and magnify the antagonistic effects of these pathways on each other.

### Importance of feedback loops and its perturbation in disease state

The positive feedback of AKTp on IRS and calcium on cAMP serves in attaining a bistable response in the regulatory elements of insulin and glucagon signalling pathways, respectively. The bistability helps in maintaining the output response in ON state even at the lower levels of input stimulus once activated. This facilitates to economise the requirements of the input stimulus by hormones (insulin and glucagon) rendering sustained output response for a short-lived input stimulus. This kind of design probes to explain the pulsatile nature of insulin secretion after the glucose stimulus, wherein an initial pulse in insulin secretion may prime the signalling response to a persistent activation due to underlying structure that produces bistability. Such an importance of pulsatile insulin secretion and the underlying bistability for efficient glucose homeostasis is also reported in the experimental observations ^16^. Disturbances in first-phase insulin pulse are reported in diabetic patients, implicating the inability to switch ON the insulin response leading to insulin resistance ^51^. Therefore, loss of the AKT positive feedback can make the system more prone to type II diabetes mellitus ^17^. The model from Zhao et al. suggested that a typical protein kinase C undergoes a bistable switch-ON and switch-OFF, under the non-linear control of insulin receptor substrate 2 (IRS2) and its disturbances causing insulin resistance ^16^. Likewise, Ca-cAMP-PKA feedforward loop separately plays an important role in the catabolic space with additional bistability showing up when Ca+ positive feedback on cAMP is knocked off, indicating lack of catabolic efficiency. Such a state may render difficulty in calorie expenditure and energy availability under acute demand and may lead to hypoglycaemic conditions during physiological stress.

Several cancer tumour cells exhibit constitutive activation of PI3K/AKT/mTOR pathway ^52^. The over expression of this pathway is essential for biosynthesis and provides a fitness advantage for the highly proliferating cells, a hallmark of cancer cells ^53^. It has been argued that the dysregulation in PI3K/AKT pathway operates between two extremes states leading to either diabetes or cancer, wherein the insulin activity is either reduced or increased, respectively ^14^. Therefore, as observed from our analysis, in the case of higher levels of plasma fatty acid (2-3 folds), the bistable response in phosphorylation state shows an anabolic response indicating higher insulin activity even at normal glucose levels (Figure 3 (iii)). Such a scenario is conducive to the proliferative state increasing the probability of cancerous phenotype, and may also promote higher synthesis and storage of fuels in the form of adipose tissues, leading to obesity. Moreover, in case of a sensitive positive feedback of AKTp on IRS would result in the increased span of bistability (See Figure S12), rendering a person higher chances of becoming obese even after consuming lesser amount of calories due to sensitive and persistent anabolic response to glucose. Supporting these hypotheses, recently it has been shown that the suppression of IRS2/Akt signalling prevents hepatic steatosis, non-alcoholic fatty liver disease (NAFLD) and liver cancer ^54^ indicating the importance of balance in the anabolic and catabolic response. Hence, our analysis provides a plausible mechanism for the increased instances of cancer in obesity^55^.

On the other hand, when selective dysfunction in either AKT positive feedback or decoupling of DAG-PKC crosstalk in a tissue like liver is considered, the system exhibits insulin resistance for increasing glucose levels (Figure 4(i) & 4(iv)). Under such a condition, due to lack of hepatic glucose absorption, plasma glucose levels may increase leading to hyperinsulinemia. The sustained increased insulin levels, then may affect the other tissues (such as adipose tissue and muscle with normal insulin sensitivity) to become more anabolic and sensitize AKT/PI3K pathway increasing the probability of cellular proliferation and tendency towards obesity^56-58^. Hence our analysis provides insights on possible mechanisms by which differential insulin resistance could be responsible for diabetes, obesity and cancer, simultaneously. Furthermore, our model predicts signalling abnormalities like progressive decrease in IRS and AKT activity along with increase in aPKC and mTOR activity with increase in body mass index (BMI) in human subjects (lean, obese & T2DM) and mouse, with diet induced obesity ^59^. These observations can easily be explained by our model where increase in FA and AA levels (in obese and T2DM) lead to respective increase in PKC, mTOR-S6K1p and subsequent inhibition of IRS and downstream kinases like AKTp.

### Modulation of bistablity in metabolic state by macronutrients

Analysis of steady state profiles of signalling proteins shows that insulin and glucagon signalling modules act antagonistically to balance, dominate or subdue each other to keep plasma glucose levels under homeostatic (balanced), mildly anabolic (post balanced diet conditions), highly anabolic (post high carbohydrate diet conditions) and highly catabolic (during exercise) states. The system keeps switching between these three steady state conditions as the level of macronutrients varies in the model. Our analysis shows that differences in the regulatory nature of macronutrients yield different patterns of metabolic responses for different combinations of these macronutrients in the plasma. For instance, at very high AA levels (>4 fold) response turns highly catabolic with fluxes in glucagon signalling module dominating over insulin signalling module fluxes. Such increased catabolic flux concomitant with higher amino acids may indicate increased ammonia production in liver. In such condition, patients may demonstrate impairments in urea synthesis that is proportional to the clinical severity of their liver disease ^60^. At very high levels of circulating AA, the catabolic signalling is dominant and tissues may not be able to build protein mass. One example of such catabolic disorder is muscle sarcopenia, reported in branched-chain amino acids (BCAA) supplementation studies^61^. Moreover, in case of moderately-high protein diet, it is well established that plasma glucose levels acutely reduce as compared to low protein diet ^62^, despite a dichotomous rise in circulating glucagon levels ^63^. Our integrated model effectively explains this phenomena as at moderately high AA levels, positive feedback of AKTp on IRS is dominant (Figure 3(i) and Figure S6) that not only would stimulate anabolic pathways like protein synthesis, but also suppresses glucagon signalling and subsequent catabolic pathways like gluconeogenesis with little effect on plasma glucose concentration under these conditions.

The effect of decreasing AA and FA as input to the system is minimal on the network fluxes, as it seems that plasma glucose levels have higher control on these hormones at basal levels. Whereas on moderately increasing fatty acid levels up to 3 folds, an anabolic action would increase lipogenesis at normal glucose levels and an increase in gluconeogenesis at subnormal glucose levels during switching OFF conditions. On increasing FA levels further up to 4 folds, catabolic state is prevalent that could inhibit lipogenesis. At high AA levels up to 3 folds and subnormal to normal glucose levels catabolic state is predominate (mimicking the starvation condition and indicative of gluconeogenesis from AA) whereas at higher glucose levels the response is anabolic conducive to protein synthesis under surplus AA levels ^64^. At higher fatty acid (4 folds onwards) and glucose levels (2 fold onwards), glucagon signalling module is shut OFF and insulin signalling module is not sufficiently activated to turn the system completely anabolic. These analysis shows that the efficiency of the insulin signalling pathway is high at the moderate levels of all these macronutrients, but it is reduced with further increasing amino acid and fatty acid levels in plasma. It was also noted that the defects in the glucagon signalling can also lead to a diabetic response despite healthy insulin signalling. Therefore, the homeostatic response is the result of these two competing pathways functioning through the regulatory network.

The dietary effect on these pathways aid in modulating the strength of these feedbacks leading to alterations in the output response of the pathway. In case of high protein & fat diet, fatty acids and amino acids have inhibitory effect on the insulin signalling pathway by increasing the serine phosphorylation of IRS via the activation of PKC and S6K respectively ^47,65^. This reduces the antagonistic effect of insulin on glucagon signalling pathway by further activation of cAMP and PKA. Since PKA is sensitive to suppression by insulin signalling and glucose, slightly lowering either of these can activate PKA to a higher value. Hence under higher fatty & amino acid conditions the catabolic activation is prevalent, which may further lead to higher output and lower intake of glucose by tissues like liver and muscle leading to a diabetic state. Our analysis indicates that an optimal composition of macronutrient exists for which the metabolic response can be maximized as per the requirement of physiological conditions.

### Future direction

Our model analysis would help in standardizing the dietary macronutrient composition under disease condition and also identifying the underlying mechanisms in certain metabolic diseases. Further, integration of this model with tissue metabolism models can help us identify strategies for disease mitigation using integrated model with metabolism. For example, exploring the potential strategies to counter obesity-linked disorders by reducing adipose tissue lipolysis to diminish the mobilisation of FAs and lower their plasma concentrations ^66^. Moreover, it would be interesting to analyse the integration of multiple bistable loops arising at metabolic ^19^ and signalling levels that provide highly versatile metabolic regulatory landscape for energetics adaptation under different combinations of plasma macronutrient concentrations and its disturbances in disease states.

## Method

### Mathematical model development

The current model integrates previously validated mechanistic models of signalling pathways relating to insulin ^17,67^, GPCR ^40^, Ca-DAG ^39^, cAMP-PKA ^68^ with empirical models of insulin and glucagon secretion ^64,69^ to generate an integrated model with 36 state variables. The integrated model contains several feedbacks and crosstalk accounting for the antagonistic nature of insulin and glucagon signalling. The ODEs have been formulated based on kinetic rate law and mass balance of signalling proteins. Hormone secretion kinetics has been quantified by accounting for the effects of plasma macronutrient levels (Supplementary material –Appendix II). Dynamic solution of these ODEs are obtained using ODE15s solver in MATLAB. The steady state profiles for signalling components are obtained and analysed for corresponding levels of glucose, fatty acids and amino acids in plasma.

There are three modules in the integrated insulin-glucagon signalling network (Figure 1) that represent both the hormones and the macronutrients (glucose, amino acids and fatty acids) in blood. Plasma concentrations of macronutrients and hormones act as input to the insulin and glucagon signalling pathways. The governing equations capturing the interplay between these pathways are given in Appendix II (supplementary material). Hill functions are used to capture important feedbacks and crosstalk in the network. Positive feedbacks of AKT on IRS, Ca on cAMP, cAMP on PKA along with negative feedbacks of S6K1p & PKC on IRS are the significant ones. Moreover, PKA degradation of PDE3 and PDE3 degradation of cAMP form a double negative feedback loop on the glucagon signalling module. Mutually antagonistic actions of both the signalling modules is modelled by the two crosstalk. Firstly, DAG activates PKC that inhibits IRS and secondly AKT activates PDE3 which promotes cAMP degradation and reduces the levels of PKA in the network ^70^. The model was qualitatively validated by matching the output simulation of the existing model with the simulations of the source models and the data from literature (see Figure S1).

### Phosphorylation state

The outputs of signalling pathways characterising insulin and glucagon signalling modules are phosphorylated AKT and activated PKA, respectively. The activated levels of AKT and PKA indicate the anabolic and catabolic state of a cell. In order to obtain the overall metabolic state of a cell, we quantify phosphorylation state metric ‘*P*_*s*_’ as a function of activated AKT and PKA levels ^71^ as follows:

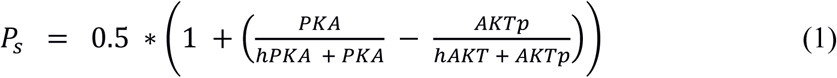

where, *h*_*pka*_ and *h*_*akt*_ are the half saturation thresholds for the signalling parameters. Figure 2(iii) shows the phosphorylation state with respect to the varying glucose input to the system. *P*_*s*_ below 0.45 depicts the anabolic zone, *P*_*s*_ within 0.45 to 0.55 depicts the homeostatic zone and above 0.55 depicts the catabolic zone.

In order to visualise the relative operational strengths of the feedbacks and crosstalk under various physiological conditions, the absolute values of the corresponding Hill functions were plotted in the network diagrams as reported in the supplementary information (Figure S2 – S11).

## Acknowledgement

We are thankful for the teaching assistantship provided by Indian Institute of Technology, Bombay through MHRD, India, to support this work.

## Author contributions

PRS and KVV conceived the work, PRS and MT performed analysis, PRS, MT and KVV wrote the manuscript.

## Supplementary Information

Appendix I-Insulin and glucagon signalling pathway

Appendix II-Details on mathematical modelling (parameters and equations)

Appendix III-Supplementary Figures

