## Supplemental Material for "Computational Analysis of Insulin-Glucagon Signalling Network: Implications of Bistability in Metabolic Homeostasis and Disease states"

### Supplementary Material

### Appendix-I

#### Insulin signaling

Insulin signalling is brought about by a complex network of synchronised events. Insulin binds to its cell surface receptor (IR) and induces a conformational change on the receptor leading to its activation [1,2]. On activation, the insulin receptor (IR) phosphorylates several members of the insulin receptor substrate family (IRS 1 & IRS 2) [3]. IRS triggers activation of PI3K and conversion by its catalytic domain of phosphatidylinositol (3,4)-bisphosphate ($PIP_2$) lipids to phosphatidylinositol (3,4,5)-trisphosphate ($PIP_3$). PKB/Akt binds to $PIP_3$ at the plasma membrane and subsequently gets activated. This PI3K-AKT pathway is highly conserved in most of the metabolic actions of insulin and is a critical node in the insulin signalling pathway [4,5]. PI3K also activates PKC [6] which negatively regulates the function of IRS proteins [7-9]. Similar findings suggest AKT to be a positive regulator of IRS activation [10,11]. Further, AKT and PKC activate GLUT4 translocation to the cell surface [12,13] subsequent to which glucose is transported inside the cell. Similarly, stimulated by insulin, fatty acid transport protein (FATP) and amino acid transport proteins (AATP) play a role in absorption of fatty acids and amino acids, respectively [14,15] and initiate the anabolic processes in cells when the concentration of these metabolites become excess in plasma. Moreover, activation of mTOR pathways is essential for protein synthesis which is regulated by insulin & amino acids [16]. Insulin regulates the activation of mTOR through the action of AKT whereas amino acids directly activate mTORc2 complex that phosphorylates S6K1 and further activates it for protein synthesis. S6K1p exerts a negative feedback on insulin signalling by serine phosphorylation of IRS1 adaptor molecule and also inhibition of formation of mTOR-rictor complex which in turn activates AKT[17,18]. Moreover, AKT also activates PDE3 which enhances degradation of cAMP mediated glucagon signalling [19].

**Glucagon signalling**

Glucagon, on the other hand, acts opposite to insulin action both at the signalling and secretion levels. It initiates the breakdown of glycogen and triglyceride stores for glucose release from muscle & liver and fatty acid release from adipose tissues, respectively, during exercise and fasting conditions [20]. Glucagon signalling mechanisms trigger metabolic pathways via second messengers like cyclic adenosine monophosphate (cAMP), cAMP dependent protein kinase (PKA) & calcium. Glucagon binds to the G protein coupled receptors (GPCR) present on the cell surface; thereby altering the conformation of the cytoplasmic domain of GPCR. Activated GPCR binds to trimeric GTP binding proteins (G proteins) via GDP-GTP exchange which relays the intracellular signal downstream to G-protein linked receptors [21-24]. Gs and Gq are the two different kinds of G proteins associated with the glucagon receptors which relay two distinct signalling pathways [25]. Gs protein activate the adenylate cyclase enzyme which leads to the increase in cAMP concentration that binds and activates PKA in the cells. Gq proteins activate PLC that activates the secondary messenger IP3, which in turn stimulates the release of intracellular calcium from the endoplasmic reticulum stores. Both activation of glycogen phosphorylase and increase in calcium levels promote glycogen breakdown in muscle and liver [20,26]. PLC hydrolyses the membrane phospholipid $PIP_2$ to form $IP_3$ and diacylglycerol (DAG). DAG recruits PKC attaching it to the plasma membrane, whereas $IP_3$ diffuses to the ER and binds to an $IP_3$ receptor that serves as a $Ca^{2+}$ channel and releases calcium ions from the ER into the cytosol. PKC interacts with DAG via its C1 domain by undergoing a conformational change, thus enabling it to phosphorylate one of its substrate IRS and thereby inhibiting insulin signalling and subsequent glucose uptake [27].

### Appendix II

### Mathematical model for Insulin-glucagon signaling network

| Parameter | Value | Units | Parameter | Value | Units |
|---|---|---|---|---|---|
| nakt | 1.8 | - | Glcn_recp | 9*1E-13 | M |
| npkc | 1 | - | a4 | 6*1E7 | /(M.min) |
| kakt | 4.25 | % | a3 | 0.1 | /min |
| kn | 7 | % | Rt | 126500 | - |
| k1aBasic | 137775 | /min | alf | 10000 | - |
| k1f | 673.445*E10 | /min | V | 11 | L |
| k1b | 543752 | /min | Vp | 3 | L |
| k2f | 141.254 | /min | c1 | 10*60 | /min |
| k21f | 9178.23*7 | /min | c2 | 100*60 | /min |
| k2b | 3533330 | /min | c3a | 312*1E -3 | /min |
| k4f | 226300 | /min | c4 | 4*60*1E-3 | /min |
| k4b | 1493960 | /min | c5 | 5.2*60*1E-3 | /min |
| k4f1 | 513844 | /min | c8 | 650*60 | /min |
| k4b1 | 320822 | /min | A0 | 3 | AU |
| k6f | 1337290/2 | /min | B1 | 100 | AU |
| k6b | 1629.22*20 | /min | B2 | 1E6 | AU |
| k5f | 0.06308 | /min | kfdag | 2*60*0.01 | /min |
| k5b | 0.212 | /min | vf | 60 | /min |
| k8f | 856353 | /min | kbdag | 0.5*60 | /min |
| k8f1 | 856353/2 | /min | kc1 | 2.0*1E -6 | µM /min |
| k8b | 1430390*1.5 | /min | kcm1 | 25*1E -12 | M |
| AMPK_eff | AMPK | AU | kc2 | 1.18 | /min |
| PP2Amax | 5 | M | kcm2 | 1 | AU |
| f_20 | 1.5E-3 | /min | pn | 1 | - |
| f_21 | 5E-4 | /min | pe | 2 | - |
| ns | 3 | - | cqi | 2 | AU |
| faa | 0.75 | mM | Va1 | 0.9 | /min |
| Vi | 0.05 | l/kg | Va2 | 8 | /min |
| m1 | 0.19 | /min | kcamp1 | 2*3.2*1E -6 | µM |
| m2 | 0.484 | /min | kcamp2 | 4*3.2*1E -6 | µM |
| m4 | 0.194 | /min | PKAt | 6*1E-4 | µM |
| m5 | 0.0304*Vi | /min | $K_{akt1}$ | 0.3 | /min |
| m6 | 0.6471 | - | kpde | 1.5 | /min |
| Gamma | 0.5 | /min | PDE3t | 5 | AU |
| AMPKt | 1 | AU | PKA_L | PKA/(8*1E-6) | - |
| kam1 | 1 | mM | ks | 1E-4 | µM |
| Kam2 | 2.25 | mM | Kbpde3 | 0.001 | /min |
| a1 | 0.12*1.25 | /min | KAMP_L | 0.16 | mM |
| a2 | 0.3 | /min | KATP_L | 2.8 | mM |
| Gm | 1.35*1E-10 | M/min | Kmpr | 2 | AU |
| p1 | 0.005*180 | /(mg/L) | BsAla | 0.25 | mM |
| q1 | 10 | - | Vala | 2.50E-10 | M |
| Vglu | 48*E-12 | M/min | Vgprt | 26 | AU/min |
| ng | 4.65 | - | klrs | 95000 | AU |
| kmg | 8.9 | mM | Vplc | 233 | AU/min |
| kma | 0.6 | mM | kgprt | 32 | AU |
| kmf | 1.6 | mM | Vip3 | 16.3 | AU/min |

| | | | | | |
|---|---|---|---|---|---|
| *na* | 5.8 | - | *kplc* | 42 | AU |
| *nf* | 4.8 | - | *Vcal* | 35 | AU/min |
| *Vaa* | 25*E-12 | M/min | *kip3* | 9 | AU |
| *Vfa* | 25*E-12 | M/min | *kgcn* | 3.20E-05 | M |
| *Vsk* | 0.2 | AU | *kmak* | 0.1 | % |
| *Kmsk* | 6 | AU | *kpka* | 12 | AU |
| *Kmfa* | 1.5 | mM | *Vakn* | 5 | AU |
| *Kmdg* | 7 | mM | *kman* | 0.05 | % |
| *Vmt* | 2.5 | AU | *kipk* | 3 | AU |
| *Vpi3* | 3 | AU | *Kmp2* | 6 | % |
| *Kmp3* | 4 | % | *Kmgln* | 25E-12 | M |
| *Vgcn* | 2 | AU | | | |

***Insulin secretion kinetics***

$$Insec = Vglu * \left(\frac{Ca_{glu}^{ng}}{Ca_{glu}^{ng}+kmg^{ng}}\right) + Vaa * \left(\frac{Ca_{ala}^{na}}{Ca_{ala}^{na}+kma^{na}}\right) + Vfa * \left(\frac{Ca_{ffa}^{nf}}{Ca_{ffa}^{nf}+kmf^{nf}}\right)$$

$$ISR = Insec/10^{-12}$$

***Insulin in portal vein***

$$\frac{d(Ins_V)}{dt} = -\,(gamma * InsV)\; +\; ISR$$

$$InSec\; =\; gamma * InsV$$

***HE-Hepatic Extraction***

$$HE\; =\; (-m5 * Insec)\; +\; m6$$

$$m3\; =\; (HE * m1)/(1 - HE)$$

***Insulin in Liver***

$$\frac{dIns_L}{dt} = -(m1 + m3) * InsL\; +\; m2 * InsP\; +\; InSec$$

***Insulin in Plasma***

$$\frac{dIns_P}{dt} = -(m2 + m4) * InsP\; +\; m1 * InsL$$

*Effect of glucagon on plasma insulin*

$$insulin_p = \frac{Ins_p * 10^{-12}}{Vi} * Vgln * \frac{Kmgln^2}{Gln_p^2 + kmgln^2}$$

$$v_{1f} = k_{1f} * insulin_p * 1000 * IR + K_{1aBasic} * IR$$

$$v_{1b} = k_{1b} * IR_p$$

$$v_{2f} = k_{2f} + k_{21f} * \left(\frac{AKT_p^{n_{akt}}}{AKT_p^{n_{akt}} + k_{akt}^{n_{akt}}}\right) * IRp * IRS * \frac{k_n}{k_n + \left(\frac{PKC_p}{1.5}\right)^{n_{pkc}}}$$

***S6K negative feedback on IRS1***

$$S6K_{Ntv} = Vsk * (\frac{S6K1a^4}{kmsk^4 + S6K1a^4})$$

$$v_{2b} = k_{2b} * IRS_p * (1 + S6K_{Ntv})$$

$$v_{4f} = k_{4f} * PI3K * IRS_p$$

$$v_{4b} = k_{4b} * PI3K_p$$

$$v_{4f1} = k_{4f1} * AKT * PI3K_p$$

$$v_{4b1} = k_{4b1} * AKT\_p$$

***Positive feedback of FFA on PKC***

$$FFA_{ptv_{pkc}} = 1 + 0.5 * \left(\frac{FFA^3}{FFA^3 + kmfa^3}\right)$$

***Positive feedback of DAG on PKC***

$$DAG_{ptv_{pkc}} = 1 + \frac{DAG^3}{DAG^3 + kmdg^3}$$

$$v_{6f} = k_{6f} * FFA_{ptv_{pkc}} * DAG_{ptv_{pkc}} * PKC * PI3K_p$$

$$v_{6b} = k_{6b} * PKC_p$$

$$v_{5f} = k_{5f} * \left(0.2 * AKT_p + 0.8 * PKC_p\right) * GLUT4_c$$

$$v_{5b} = k_{5b} * GLUT4_s$$

***mTOR Raptor activation by AA in absence of insulin***

$$AA_{mTOR_{eff}} = Vmt * \frac{AA^{ns}}{AA^{ns} + faa^{ns}}$$

$$v_{8f} = k_{8f} * mTOR * AKT_p + k_{8f1} * \left(1 + AA_{mTOR_{eff}}\right) * mTOR$$

$$v_{8f} = k_{8b} * mTORa$$

***S6K1 phosphorylation-dephosphorylation***

$$PDK1_a = Vpi3 * \frac{PI3K}{Kmp3 + PI3K}$$

$$PP2_a = PP2_{amax} * \frac{Kmp2^{1.2}}{Kmp2^{1.2} + mTORa^{1.2}}$$

$$AMPK_{Ntv_{S6K}} = \frac{0.75^2}{0.75^2 + AMPK_{eff}^2}$$

$$reg_1 = \frac{kmpr^4}{kmpr^4 + PP2_a^4}$$

$$v_{9f} = f_{20} * PDK1_a * mTOR_a * reg_1 * S6K1 * AMPK_{Ntv_{S6k}}$$

$$v_{9b} = f_{21} * PP2_a * S6K1_a$$

**ODEs: Mass balance equations**

$$\frac{d(IR)}{dt} = v_{1b} - v_{1f}$$

$$\frac{d(IR_p)}{dt} = v_{1f} - v_{1b}$$

$$\frac{d(IRS)}{dt} = v_{2b} - v_{2f}$$

$$\frac{d(IRS_p)}{dt} = v_{2f} - v_{2b}$$

$$\frac{d(PI3K)}{dt} = v_{4b} - v_{4f}$$

$$\frac{d(PI3K_p)}{dt} = v_{4f} - v_{4b}$$

$$\frac{d(AKT)}{dt} = v_{4b1} - v_{4f1}$$

$$\frac{d(AKT_p)}{dt} = v_{4f1} - v_{4b1}$$

$$\frac{d(PKC)}{dt} = v_{6\mathrm{b}} - v_{6\mathrm{f}}$$

$$\frac{d(PKC_p)}{dt} = v_{6f} - v_{6b}$$

$$\frac{d(GLUT4_c)}{dt} = v_{5b} - v_{5f}$$

$$\frac{d(GLUT4_s)}{dt} = v_{5f} - v_{5b}$$

$$\frac{d(mTOR)}{dt} = v_{8b} - v_{8f}$$

$$\frac{d(mTOR_a)}{dt} = v_{8f} - v_{8b}$$

$$\frac{d(S6K1)}{dt} = v_{9b} - \mathrm{v}_{9f}$$

$$\frac{d(S6K1_a)}{dt} = v_{9f} - v_{9b}$$

***Glucagon Secretion and kinetics***

$$if\ (Ca_{Ala} - bs_{Ala}) >= 0\ ;\ AmGlcn = (Ca_{Ala} - bs_{Ala});\ else\ AmGlcn = 0$$

$$AA_{Glcn_{eff}} = Vala * \left(\frac{Am_{Glcn}^{4.5}}{Am_{Glcn}^{4.5}+1}\right);\quad if\,Ca\_Glu <= 5;$$

$$Glcn_{Sec} = \frac{Gm}{\frac{Ca_{Glu}}{5} + q_1 * \exp(p_1 * (Ca_{Glu} - 5))} + AA_{Glcn_{eff}};\ else$$

$$Glcn_{Sec} = \frac{Gm}{1+q_1*\exp(p_1*(Ca_{Glu}-5))} + AA_{Glcn_{eff}}$$

***Plasma Glucagon conc.***

$$\frac{dGln_p}{dt} = -(a1 + a2) * Gln_p + Glcn_{Sec}$$

***Glucagon receptor model***

$$GPRT = Vgprt * \frac{LRS}{klrs + LRS}$$

$$plc = Vplc * \frac{GPRT^2}{kgprt^2 + GPRT^2}$$

$$IP3i = Vip3 * \frac{plc^4}{kplc^4 + plc^4}$$

$$Cal = Vcal * \frac{IP3i^{1.8}}{kip3^{1.8} + IP3i^{1.8}}$$

$$\frac{dGlcn}{dt} = a_1 * V_p * \frac{Glnp}{V} - a_3 * Glcn - a_4 * Glcn * \frac{FR}{Rt} * Gln_{recp}$$

$$GCN = Glcn * 10^6$$

$$c_3 = c_{3a} * Vgcn * \frac{GCN}{kgcn + GCN}$$

***FR-free receptor***

$$\frac{dFR}{dt} = c_1 * LR - c_2 * GCN * FR - c_3 * FR + c_4 * RS$$

***RS-Sequesterd receptor***

$$\frac{dRS}{dt} = c_1 * LRS - alf * c_2 * GCN * RS + c_3 * FR - c_4 * RS$$

***LR-Ligand bound receptor***

$$\frac{dLR}{dt} = c_2 * GCN * FR - c_1 * LR + \frac{c_4}{alf} * LRS - c_3 * LR + c_5 * LRP$$

***LRS-ligand bound sequestered receptor***

$$\frac{dLRS}{dt} = c_3 * LR - \frac{c_4}{alf} * LRS + alf * c_2 * GCN * RS - c_1 * LRS - c_8 * \left(1 + A_0 * \frac{GPRT}{B_1 + GPRT} * \frac{LRS}{B_2 + LRS}\right)$$

***LRP***

$$\frac{dLRP}{dt} = c_8 * \left(1 + A_0 * \frac{GPRT}{B1 + GPRT} * \frac{LRS}{B_2 + LRS}\right) - c_5 * LRP$$

***DAG***

$$Kc2p = 1 - \left(\frac{PKA^4}{ks^4 + PKA^4}\right)$$

$$\frac{dDAG}{dt} = kfdag * Cal * \frac{plc}{K_{c2p} + 0.1 * Cal} + V_f * FA - kbdag * DAG$$

***cAMP-PKA signalling***

$$Kck = 1 + 0.5 * cqi * \frac{Cal^3}{9^3 + Cal^3}$$

$$Jg12 = Kck * K_{c1} * \left(\frac{Glnp^2}{K_{cm1}^2 + Glnp^2}\right) - kc2 * \frac{PDE3_a}{K_{cm2} + PDE3_a} * cAMP$$

$$Jg13 = V_{a1} * \frac{cAMP^3}{(Kcamp_1)^3 + cAMP^3} * (PKA_t - PKA) - V_{a2} * PKA * \frac{Kcamp_2}{Kcamp_2 + cAMP}$$

***PKA***

$$\frac{dPKA}{dt} = Jg13$$

***cAMP***

$$\frac{dcAMP}{dt} = Jg12 - 2 * \frac{dPKA}{dt}$$

***PDE3a***

$$AKT_{ptv_{PDE3}} = K_{akt1} * \frac{AKT_p^2}{kmak^2 + AKT_p^2}$$

$$\frac{dPDE3}{dt} = AKT_{ptv_{PDE3}} * (PDE3_t - PDE3_a) - k_{pde} * \frac{PKA_L}{kpka + PKA_L} * PDE3a$$

***AMPK***

$$AKT_{ntv_{AMPK}} = Vakn * \frac{AKT_p^2}{AKT_p^2 + kman^2}$$

$$PKA_{ntv_{AMPK}} = \frac{kipk^2}{kipk^2 + PKA_L^2}$$

$$\frac{dAMPK}{dt} = k_{am1} * AMP_{ATP_{AMPK}} * PKA_{ntv_{AMPK}} * (AMPK_t - AMPK) - K_{am2} * AMPK * AKT_{ntv_{AMPK}}$$

### Appendix –III

### FIGURES

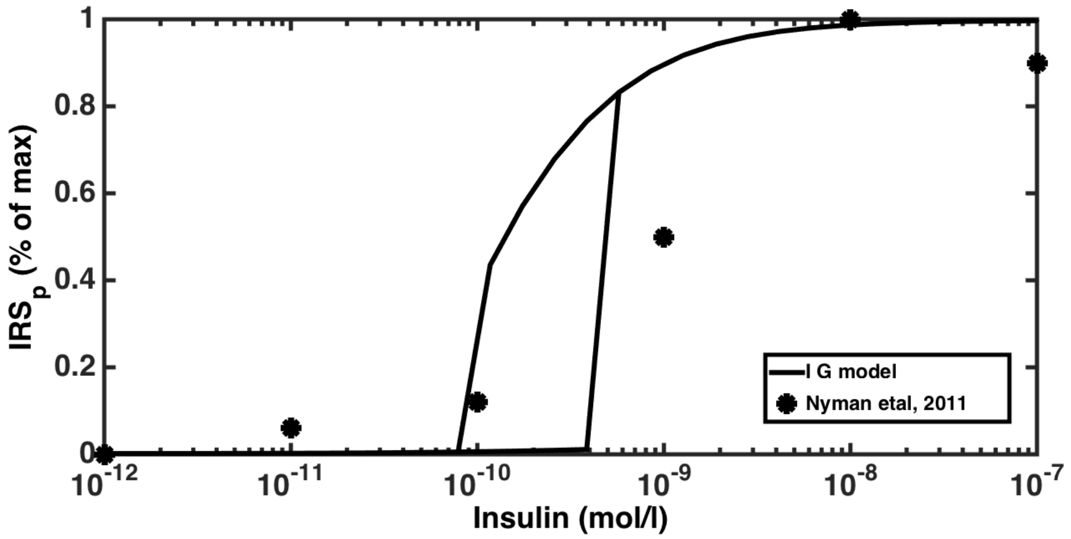


(a)

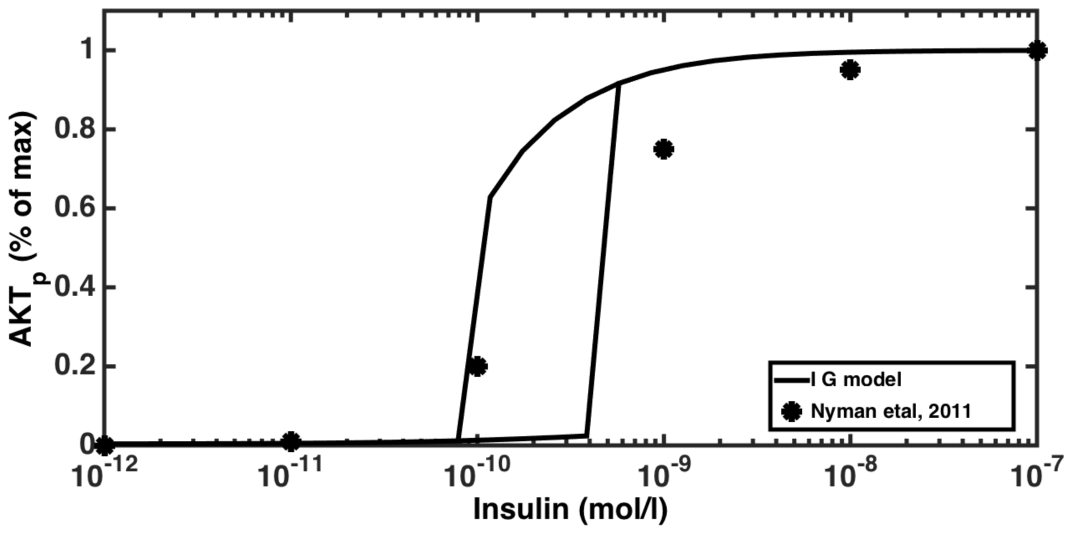


(b)

Figure S1. Model Validation. Dose response to changing concentration of insulin. Solid graph is simulated by our model which shows bistable response for increasing and decreasing levels of insulin. Black filled stars indicate experimental data obtained from the input response to isolated human adipocytes[28] for (a) phosphorelated IRS (b) phosphorelated AKT.

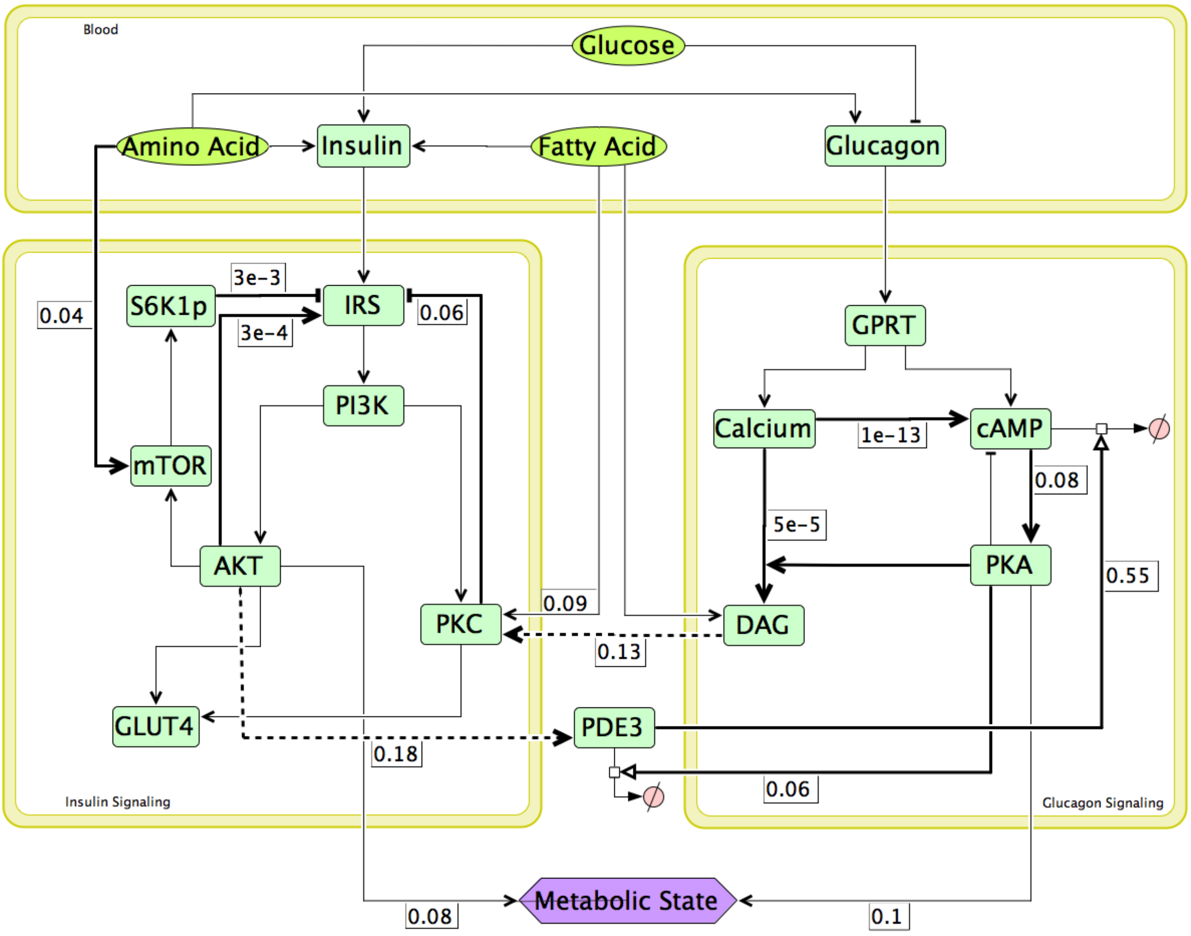


Figure S2. Network map at glucose = 1 fold, AA & FA = 1 fold i.e., during resting conditions. Numbers along the bold lines indicate magnitude of activation or inhibition in terms of the corresponding Hill function values, which lie between 0 and 1. Here most feedbacks & crosstalks are almost inactive except cAMP degradation by PDE3. Overall metabolic state is slightly catabolic with glucagon signalling module being more active than insulin signalling module (AKT contribution = 0.08 and PKA contribution = 0.1).

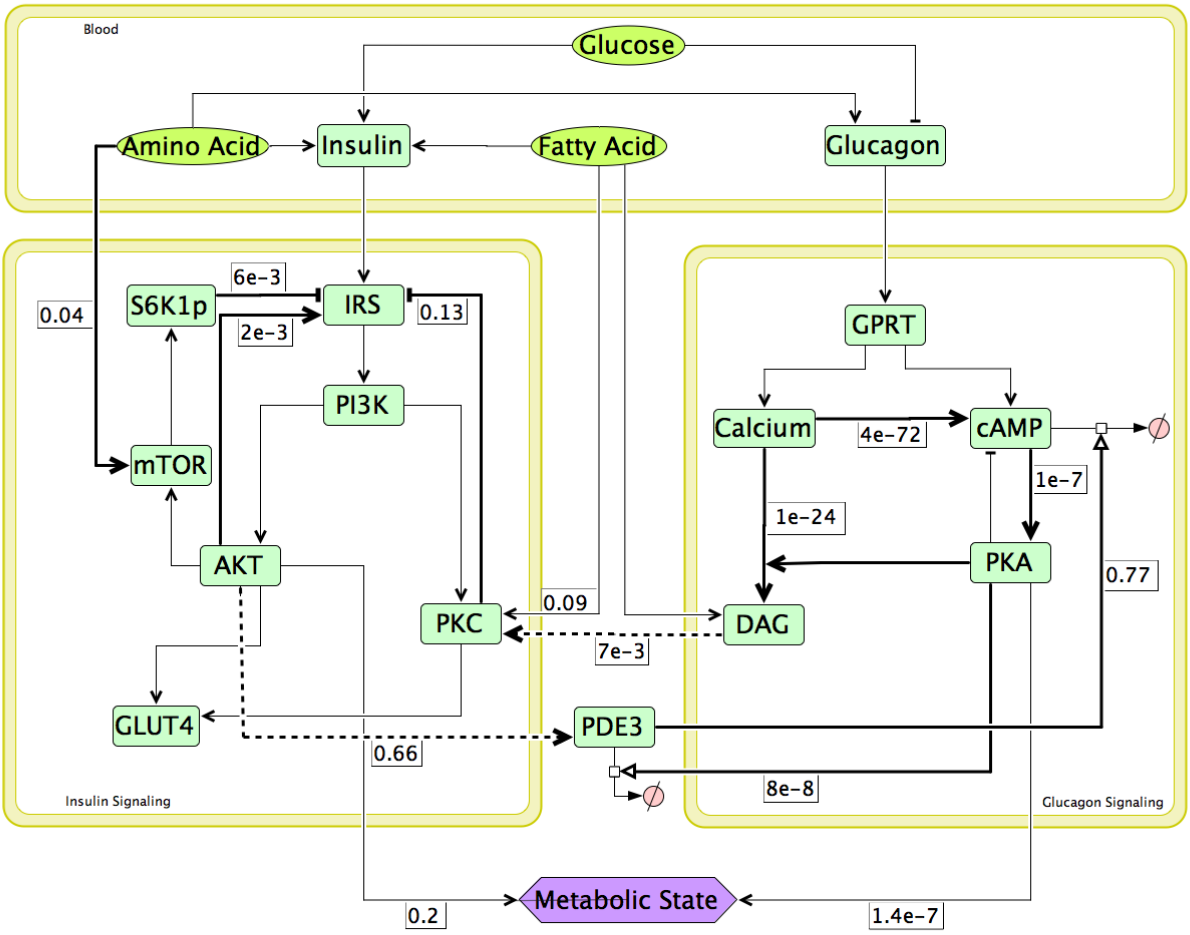


Figure S3. Network map at glucose = 1.5 fold, AA & FA = 1 fold during switching ON conditions. Numbers along the bold lines indicate magnitude of activation or inhibition in terms of the corresponding Hill function values, which lie between 0 and 1. Here AKT activation of IRS has increased slightly ( = 2e-3 vs 3e-4 in Figure S1), whereas PDE3 degradation of cAMP has increased ( = 0.77 vs 0.55 in Figure S1) as compared to the resting state. Overall metabolic state is mildly anabolic with contribution from insulin signalling being more than glucagon signalling to the net metabolic state (AKT contribution = 0.2 and PKA contribution = 1.4e-7).

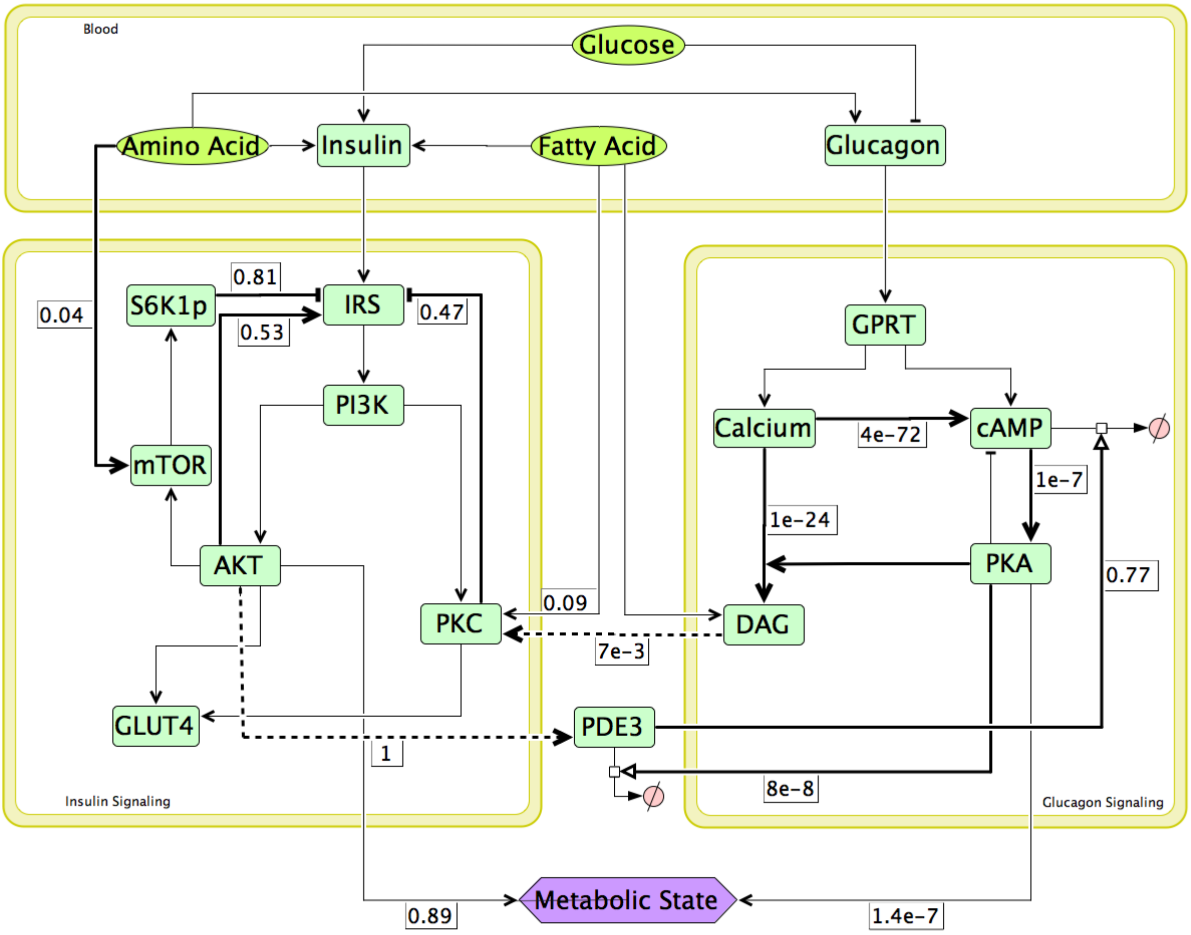


Figure S4. Network map at glucose = 1.5 fold, AA & FA = 1 fold during switching OFF conditions. Numbers along the bold lines indicate magnitude of activation or inhibition in terms of the corresponding Hill function values, which lie between 0 and 1. Here AKT positive feedback on IRS has strengthened (= 0.53 vs 3e-4 in Figure S1) along with the negative feedbacks of S6K1p (0.81 vs 3e-3 in Figure S1), PKC on IRS (0.47 vs 0.06 in Figure S1) and AKT crosstalk PDE3 (= 1 vs 0.18 in Figure S1) compared to the resting state. Overall metabolic state is heavily anabolic with contribution from insulin signalling being much more than glucagon signalling to the net metabolic state (AKT contribution = 0.89 and PKA contribution = 4.3e-8).

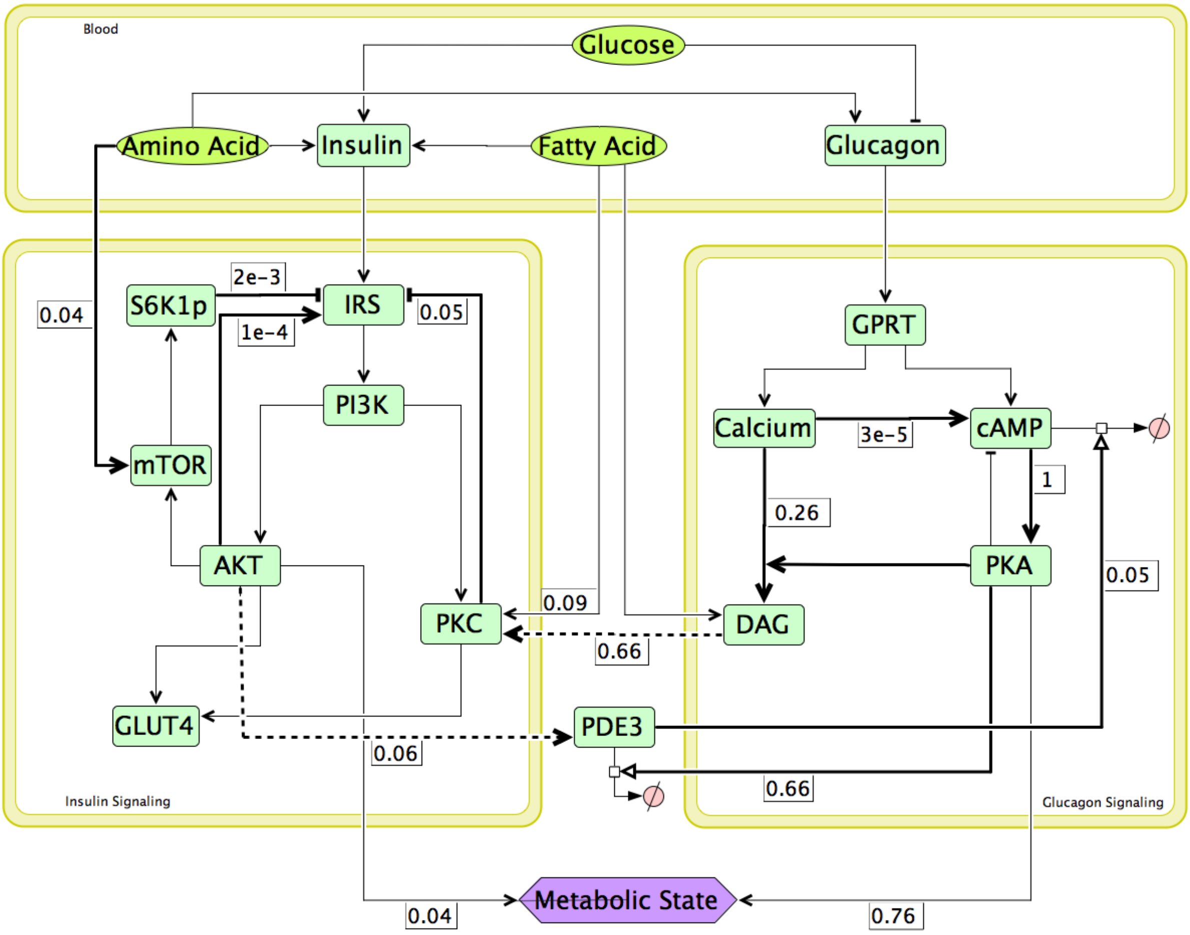


Figure S5. Network map at glucose = 0.8 fold, AA & FA = 1 fold. Numbers along the bold lines indicate magnitude of activation or inhibition in terms of the corresponding Hill function values, which lie between 0 and 1. Here PDE3 degradation of cAMP is minimal (= 0.05 vs 0.55 in figure S1), cAMP activation of PKA is maximum (=1) in this case. Hence, overall metabolic state is heavily catabolic with contribution from insulin signalling being much less than glucagon signalling (AKT contribution = 0.04 and PKA contribution = 0.76).

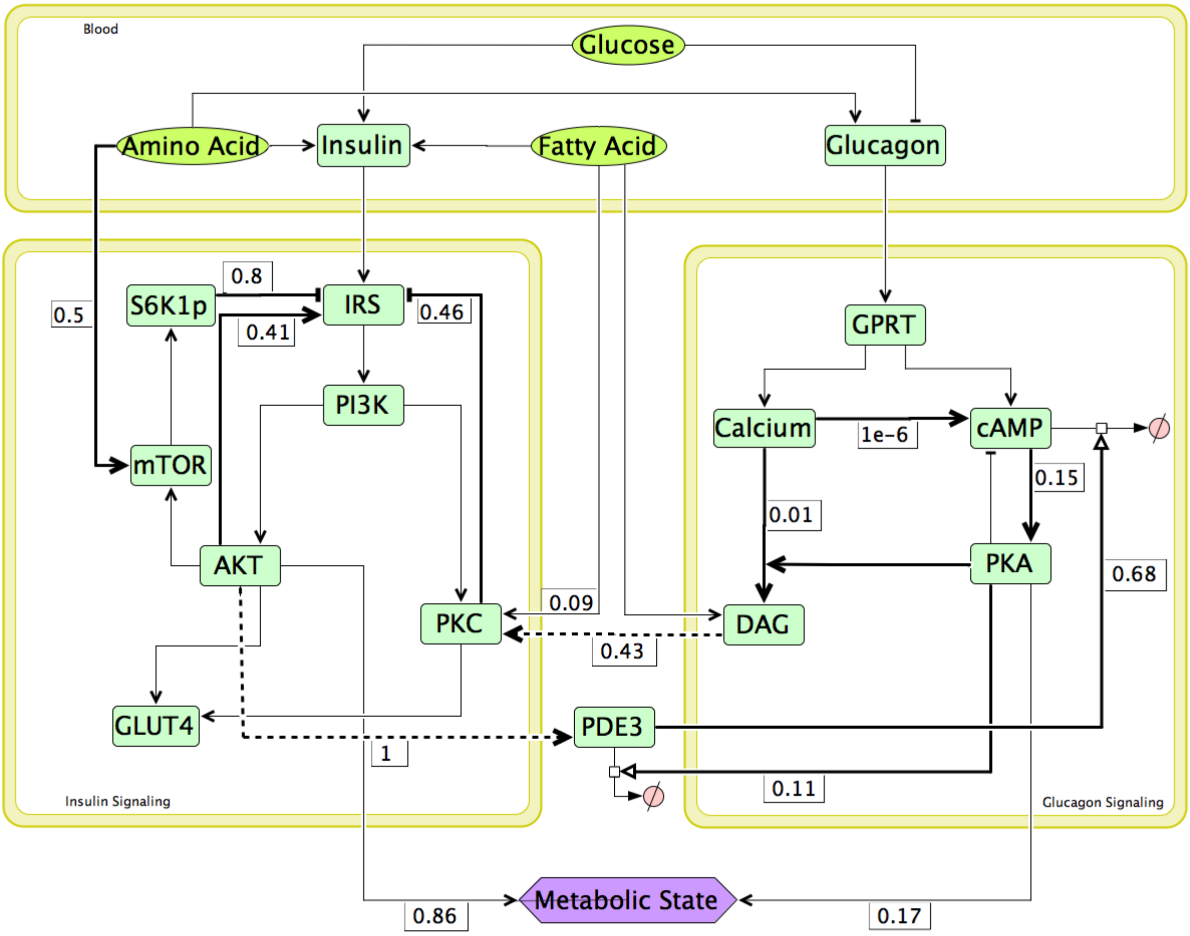


Figure S6. Network map for high plasma AA levels; with AA = 3 folds, glucose = 1 fold & FA = 1 fold for switching OFF conditions. Numbers along the bold lines indicate magnitude of activation or inhibition in terms of the corresponding Hill function values, which lie between 0 and 1. Although high AA levels are inhibiting insulin signalling pathways through increased S6K1p inhibition of IRS (= 0.8), it is also activating IRS-PI3K-AKT pathways (via increasing insulin secretion from pancreas). Moreover, AKTp positive feedback on IRS (= 0.41) and AKT crosstalk PDE3 (= 1) and cAMP degradation (=0.68) are active. Although DAG inhibition on IRS (via PKC) has increased (PKC inhibition on IRS = 0.46 vs 0.06 under resting conditions), it is not able to counter the activating effect of AKT positive feedback on IRS. Hence, overall metabolic state is heavily anabolic with contribution from insulin signalling being much more than glucagon signalling to the net metabolic state (AKT contribution = 0.86 and PKA contribution = 0.05).

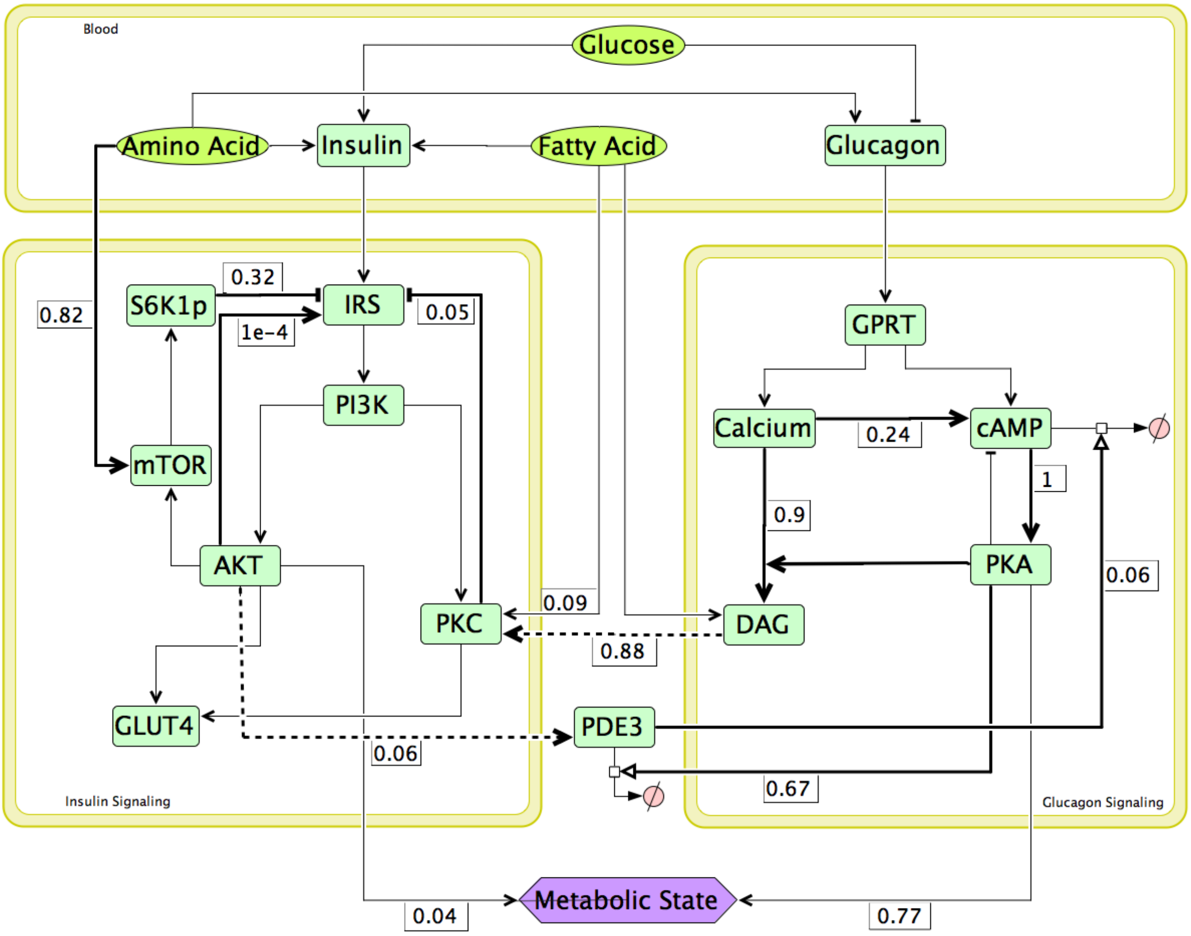


Figure S7. Network map for excess AA and high glucose levels in plasma; with glucose = 1.5 folds, AA = 5 folds & FA = 1 fold. Numbers along the bold lines indicate magnitude of activation or inhibition in terms of the corresponding Hill function values, which lie between 0 and 1. Here activation of mTOR-S6K1p pathway by plasma amino acid is inhibiting IRS as seen from the high strength of AA-mTOR activation (0.82) and S6K1p inhibition IRS (0.32). Whereas Glucagon signalling fluxes are highly activated by high amino acid levels. This leads to Ca-DAG (= 0.9) and cAMP-PKA (= 1) activations becoming very high inspite of inhibitions from high glucose levels in these condition. Insulin signalling module is almost shut off and very low levels of downstream AKTp is unable to degrade cAMP via PDE3 activation (=0.06). Hence, overall metabolic state is heavily catabolic with contribution from insulin signalling being much less than glucagon signalling to the net metabolic state (AKT contribution = 0.04 and PKA contribution = 0.77).

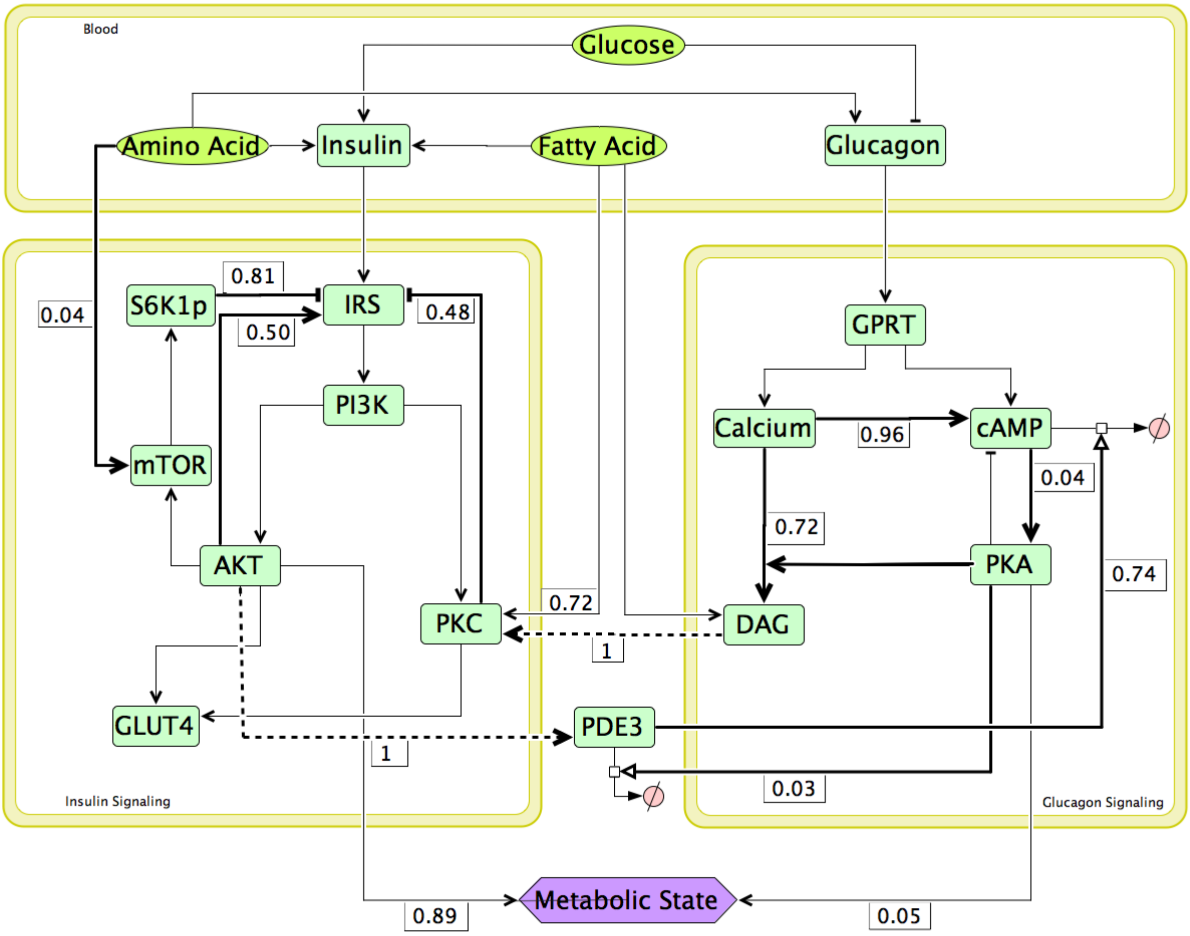


Figure S8. Network map for high FA levels in plasma; with FA = 3 folds, glucose = 1 fold & AA = 1 fold, during switching OFF conditions. Numbers along the bold lines indicate magnitude of activation or inhibition in terms of the corresponding Hill function values, which lie between 0 and 1. Here AKTp positive feedback on IRS ( = 0.50), AKT crosstalk PDE3 (=1) and PDE3 degradation of cAMP (= 0.74) are dominant like in Figure S5. Although increased AKT levels are also increasing S6K1p inhibition of IRS (=0.81) and high FA level is also strengthening PKC inhibition on IRS (=0.48), they are not able to inhibit Insulin signalling pathways due to the dominant AKT-IRS-PI3K positive feedback loop. Contribution of AKT (=0.89) is more than PKA (0.05) to the overall metabolic state, making it highly anabolic.

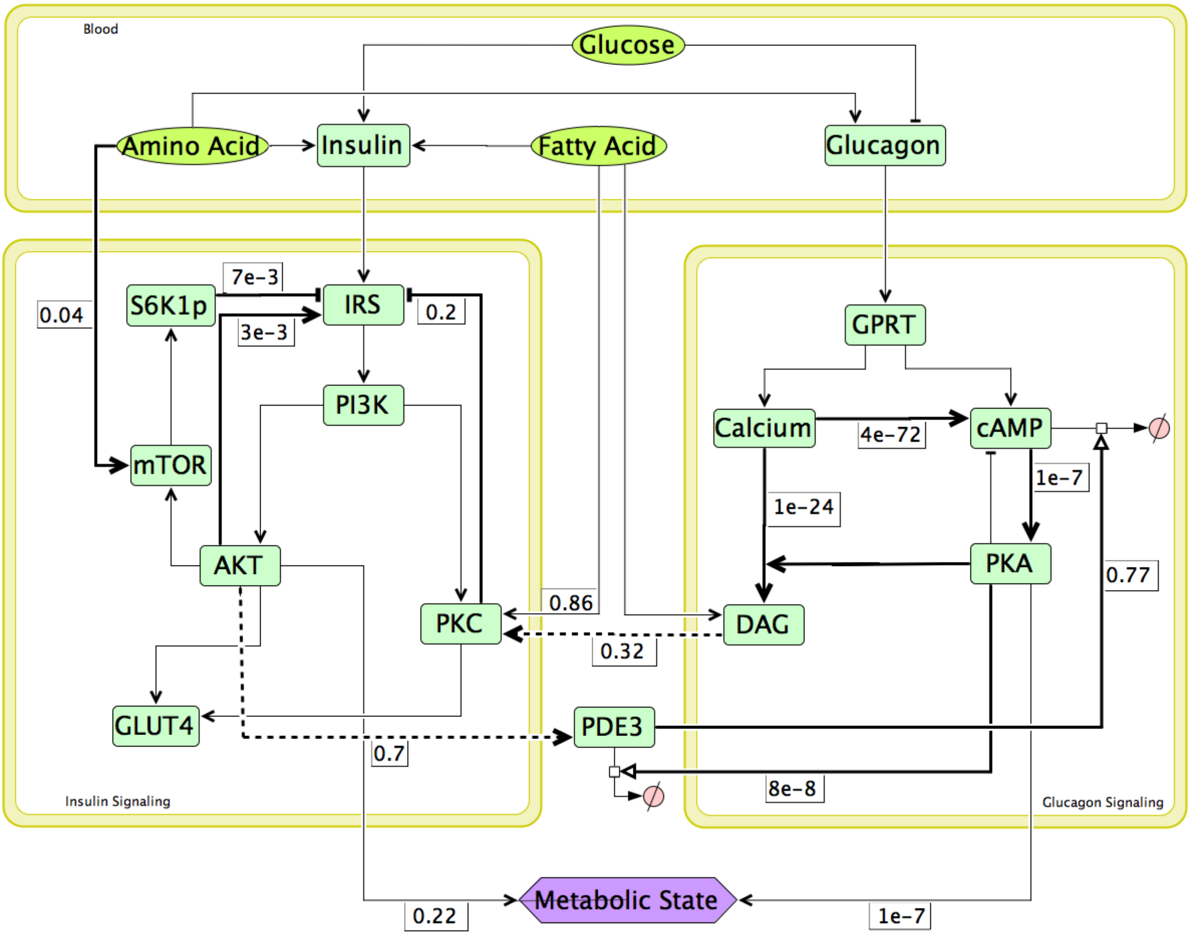


Figure S9. Network map for high FA and high glucose levels in plasma; with glucose = 1.5 folds, AA = 1 fold & FA = 4 folds. Numbers along the bold lines indicate magnitude of activation or inhibition in terms of the corresponding Hill function values, which lie between 0 and 1. Here glucagon signalling module is almost inactive due to inhibition by glucose and highly active cAMP degradation by AKTp via PDE3 (=0.77). Insulin signalling fluxes are only slightly active (inspite of increased insulin secretion by higher glucose and fatty acid levels) due to inhibition by high FA levels via PKC activation (=0.86 vs 0.09 in Figure S5). Moreover inactivated positive feedback of AKT on IRS (=3e-3) is unable to strengthen the anabolic response. Hence, overall metabolic state is only slightly anabolic with contribution from insulin signalling being more than glucagon signalling to the net metabolic state (AKT contribution = 0.22 and PKA contribution = 1e-7).

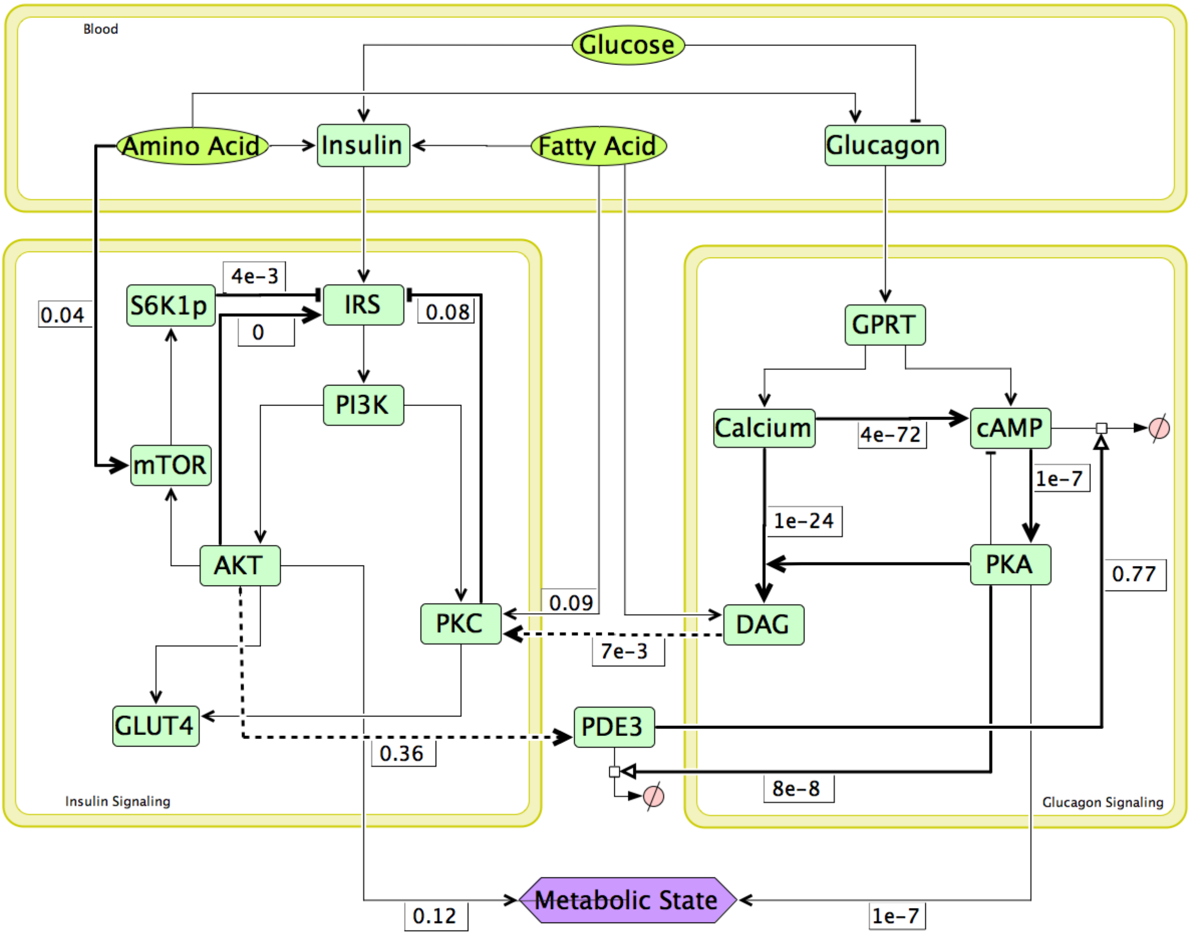


Figure S10. Network map for knockout of AKTp positive feedback on IRS; with glucose = 1.5 folds, AA = 1 fold & FA = 1 fold. Numbers along the bold lines indicate magnitude of activation or inhibition in terms of the corresponding Hill function values, which lie between 0 and 1. Glucagon signalling module is inactive due to inhibition by glucose and PDE3 degradation of cAMP (=0.77). Insulin signalling module is only slightly active (as AKTp positive feedback on IRS is knocked off). Contribution of AKT (=0.12) is more than PKA (1e-7) to the overall metabolic state, making it slightly anabolic.

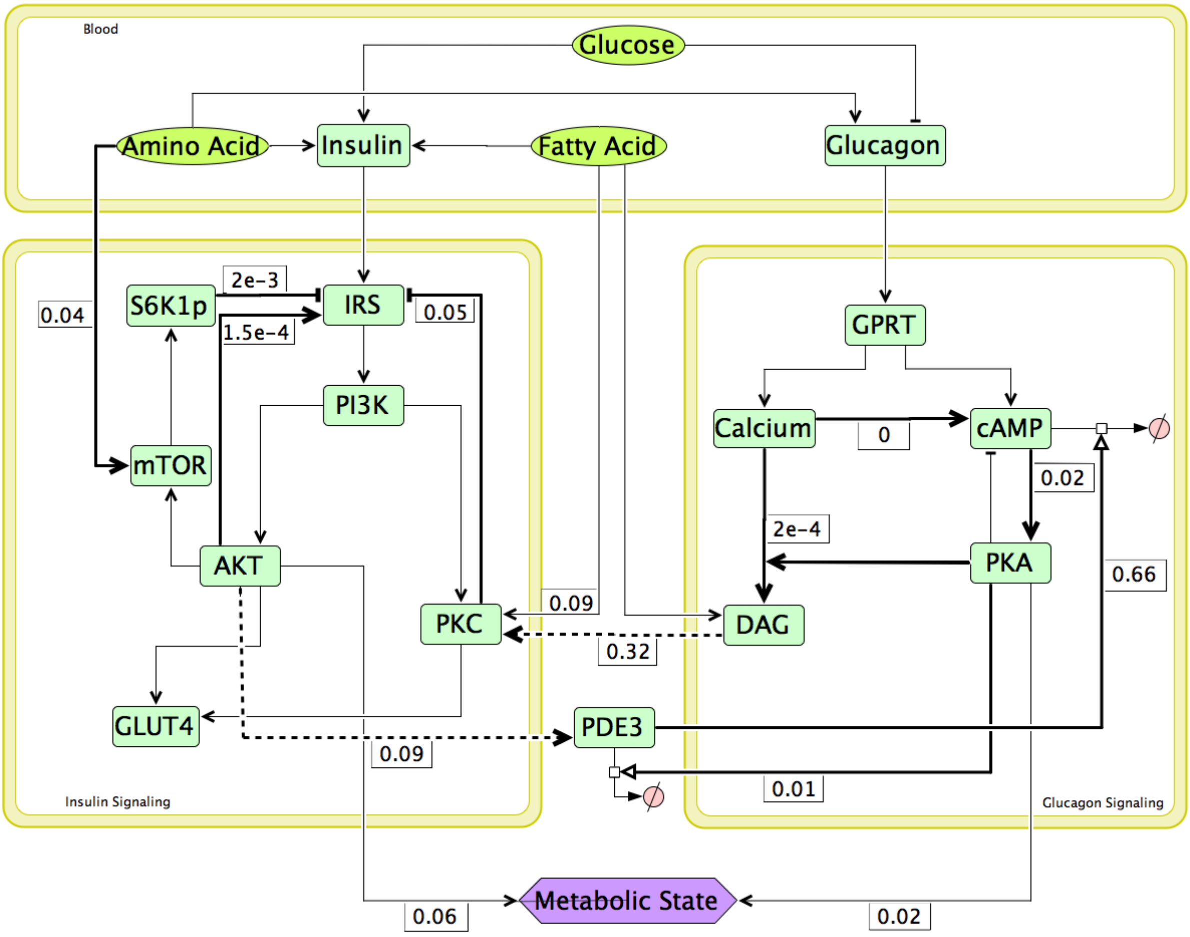


Figure S11. Network map for knockout of Ca positive feedback on cAMP; with glucose = 0.9 folds, AA = 1 fold & FA = 1 fold. Numbers along the bold lines indicate magnitude of activation or inhibition in terms of the corresponding Hill function values, which lie between 0 and 1. Here glucagon signalling module and insulin signalling module are both almost inactive leading to homeostatic conditions even at subnormal glucose levels. Knocking off positive feedback of calcium on cAMP and PDE3 degradation of cAMP (=0.66) are deactivating glucagon signalling fluxes. Contribution of both AKT (=0.06) and PKA (0.02) are minimal to the overall metabolic state, making the response homeostatic.

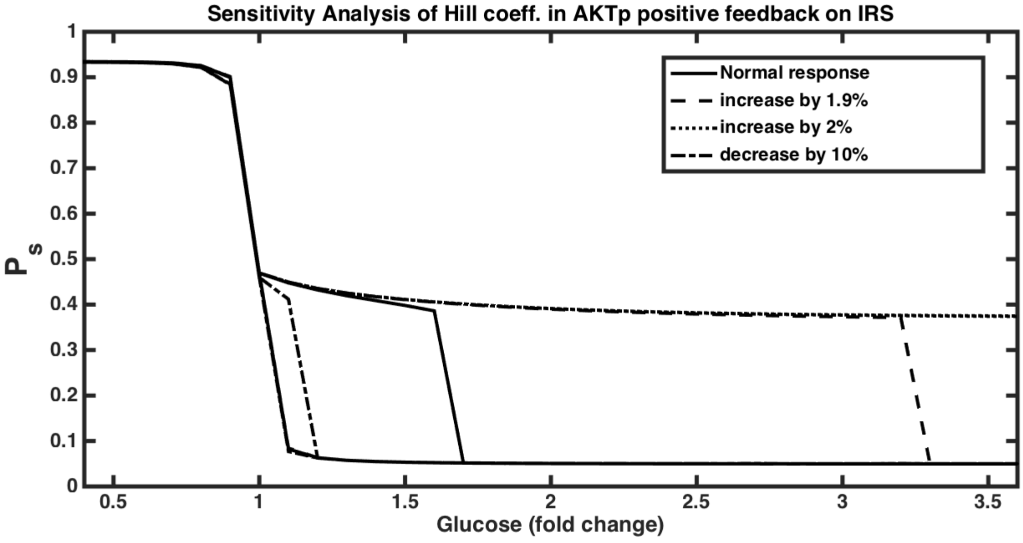


(a)

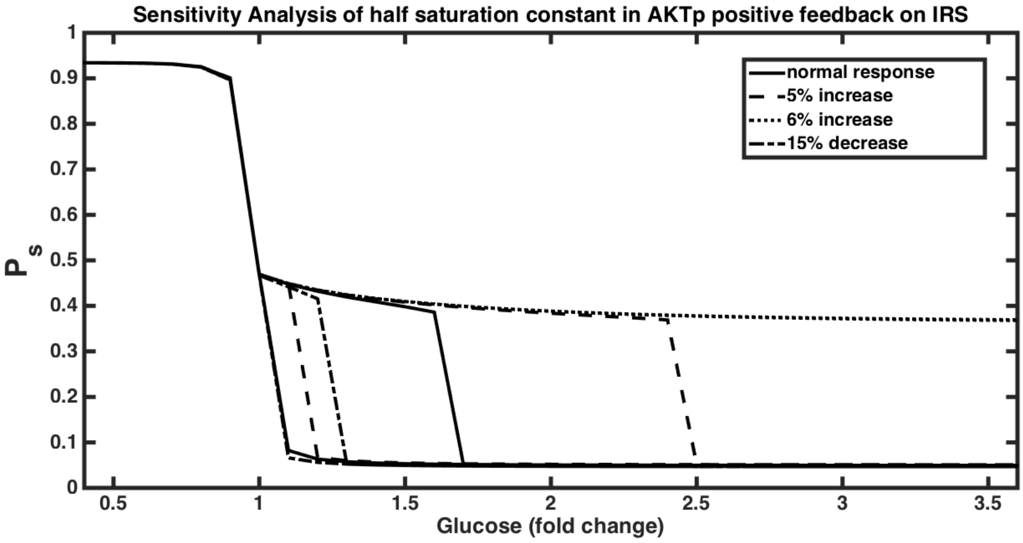


(b)

Figure S12. Sensitivity Analysis of AKTp positive feedback on IRS. (a) As the value of hill coefficient is increased up to 1.9%, bistability span increases in the anabolic space. Beyond 2% increase response turns monostable and homeostatic. While on decreasing, bistability span decreases up to 10% and beyond that it turns monostable. (b) As the value of half saturation constant is increased up to 5%, bistability span increases in the anabolic space. Beyond 6% increase response turns monostable and homeostatic. While on decreasing, bistability span decreases upto 15% and beyond that it turns monostable.

Given the importance of this feedback on the bistable response we have also performed sensitivity analysis of the hill function representing the positive feedback of AKTp on IRS (Supplementary Figure 12 (a) & (b)). For the hill coefficient parameter n, bistable response is maintained for changes ranging from -20% to +1.8% and for the half saturation constant k, bistable response is maintained for changes ranging from -15% to +10%. As the values of n & k are increased, bistability span increased and response turned monostable beyond increase in n = 1.8% & k = 10%. Whereas, as the values of n and k are reduced, bistability span decreases in the anabolic space for n up to 20% and k up to 15% reduction.
